# Precise visuomotor transformations underlying collective behavior in larval zebrafish

**DOI:** 10.1101/2021.05.24.445521

**Authors:** Roy Harpaz, Minh Nguyet Nguyen, Armin Bahl, Florian Engert

**Affiliations:** Department of Molecular and Cellular Biology, Harvard University, Cambridge 02138, USA; Center for Brain Science, Harvard University, Cambridge, Massachusetts 02138, USA; Swiss Federal Institute of Technology Lausanne (EPFL), 1015 Lausanne, Switzerland; Centre for the Advanced Study of Collective Behaviour, University of Konstanz, Konstanz 78464, Germany

## Abstract

Complex schooling behaviors result from local interactions among individuals. Yet, how sensory signals from neighbors are analyzed in the visuomotor stream of animals is poorly understood. Here, we studied aggregation behavior in larval zebrafish and found that over development larvae transition from overdispersed groups to tight shoals. Using a virtual reality assay, we characterized the algorithms fish use to transform visual inputs from neighbors into movement decisions. We found that young larvae turn away from retinal “clutter” by integrating and averaging retina-wide visual inputs within each eye, and by using a winner-take-all strategy for binocular integration. As fish mature, their responses expand to include attraction to low retinal clutter, that is based on similar algorithms of visual integration. Using model simulations, we show that the observed algorithms accurately predict group structure over development. These findings allow us to make testable predictions regarding the neuronal circuits underlying collective behavior in zebrafish.

## Introduction

Complex collective behaviors such as schooling in fish and flocking in birds can result from local interactions between individuals in the group (1–6). Understanding how sensory signals coming from surrounding neighbors guide fast and accurate movement decisions is therefore central to the understanding of emergent collective behavior and is of great interest from both computational and neurobiological perspectives (7).

Theoretical models treating groups of animals as physical systems, suggested ‘simple’ interactions among individuals and explored the collective group states that emerge from these interactions (2, 3, 5, 6, 8–12). Experimental studies, relying on recent advances in machine vision and tracking algorithms (13–18), attempted to infer individual interaction rules directly from animal movement trajectories, and compared them to the hypothesized rules from theoretical studies (19–26). Commonly, such interaction rules assume that an individual animal classifies all neighbors as individual objects, and that various computations are subsequently performed on these objects. These computations include estimating every neighbor’s distance, orientation or velocity and performing mathematical operations such as averaging and counting on these representations (2–6), or to selectively respond to specific neighbors but not to others (19, 20, 23). Alternatively, complex collective behaviors can also emerge from more simplified computations that rely primarily on the spatial and temporal low-level statistics of retinal inputs (10, 27, 28). Specifically, several theoretical models have used the visual projection of neighbors on the retina as the sole input to the animal and explored the resulting collective behavior (10, 27, 28). Whether or not animals use representations of their neighbors as individual objects, and perform complex computation on these representations or whether they base their behavioral decisions on global sensory inputs is currently unknown in most animal species. Consequently, the brain mechanisms and neurobiological circuits involved in collective social behavior are mostly unknown as well.

The zebrafish model system is uniquely situated to help address this gap of knowledge. First, this fish species exhibits complex social behaviors, even at the larval stage, that are expected to have a strong visual component ((20, 24, 26, 29–31), but see (32, 33) for other modalities). Second, previous studies in larval zebrafish have successfully characterized the underlying computations and brain mechanisms of other complex behaviors such as prey capture (34–38), predator avoidance (39, 40), motor learning (41) and decision making (42–44). In many of these studies, virtual reality (VR) assays were used to systematically probe and analyze the behavioral responses of the fish, and recently, VR assays were also shown to successfully elicit social response in larval and juvenile zebrafish (31, 45, 46). Third, the zebrafish is genetically accessible (47) and, at the larval stage, can be studied using various imaging and neural activity recording techniques (48–52). Recently, new insights into the molecular pathways involved in social and collective behavior have started to emerge, detecting unique genes and neuropeptides associated with social behavior and the detection of conspecifics (53–56). Therefore, the larval zebrafish can be used to study the specific visuomotor transformation involved in collective behavior as they emerge during development, and the neurobiological circuits at their basis.

We analyzed here the collective swimming behavior in groups of larvae at different developmental stages. We detect complex group structure already at 7 days post fertilization (dpf), that strongly depends on visual information and continues to develop as fish mature. We then utilized a virtual reality assay (31, 45, 46) to vary the static and dynamical features of naturalistic as well as artificial stimulus patterns, and tested the effects of varying the statistics of these patterns on the movement decisions of the fish. Using this assay, we characterized the precise visuomotor transformations that control individual movement decisions and the interaction rules that allow fish to integrate information from multiple neighbors. Studying these transformations over development allowed us to hypothesize which of these computations are already mature in the younger larvae, and which computations continue to evolve over development. Using model simulations we verified that the identified visuomotor transformations can accurately account for the observed collective swimming behavior of groups. Finally, we used our findings to formulate predictions about the structure and function of the neural circuits that are involved in transforming visual input into movement decisions.

## Results

### Group structure in larval zebrafish depends on visual social interactions

To understand how social interactions shape group structure over development, we studied collective swimming behavior in groups of 5 or 10 larval zebrafish at the ages 7, 14 and 21 dpf, swimming freely in closed arenas. (Fig. 1A, Movies 1-3, Methods). We find that already at 7 dpf, larval zebrafish respond to their neighbors, with groups exhibiting increased dispersion compared to chance levels (Fig. 1B-C, Fig. S1A, Methods). Group structure completely disappeared when fish were tested in total darkness, confirming the strong visual component of the interactions (Fig. 1B-C). As fish matured, this repulsive tendency reversed and fish swam towards their neighbors, resulting in an age dependent increase in group cohesion, as reported previously (20, 30, 31)(Fig. 1D). Average swimming speed and alignment between fish also increased over development, while bout rate decreased (Fig. S1B-D). Among these developmental changes in behavior, we focus here on the aggregation behavior of the fish and its unique developmental trajectory.

**Figure 1.**
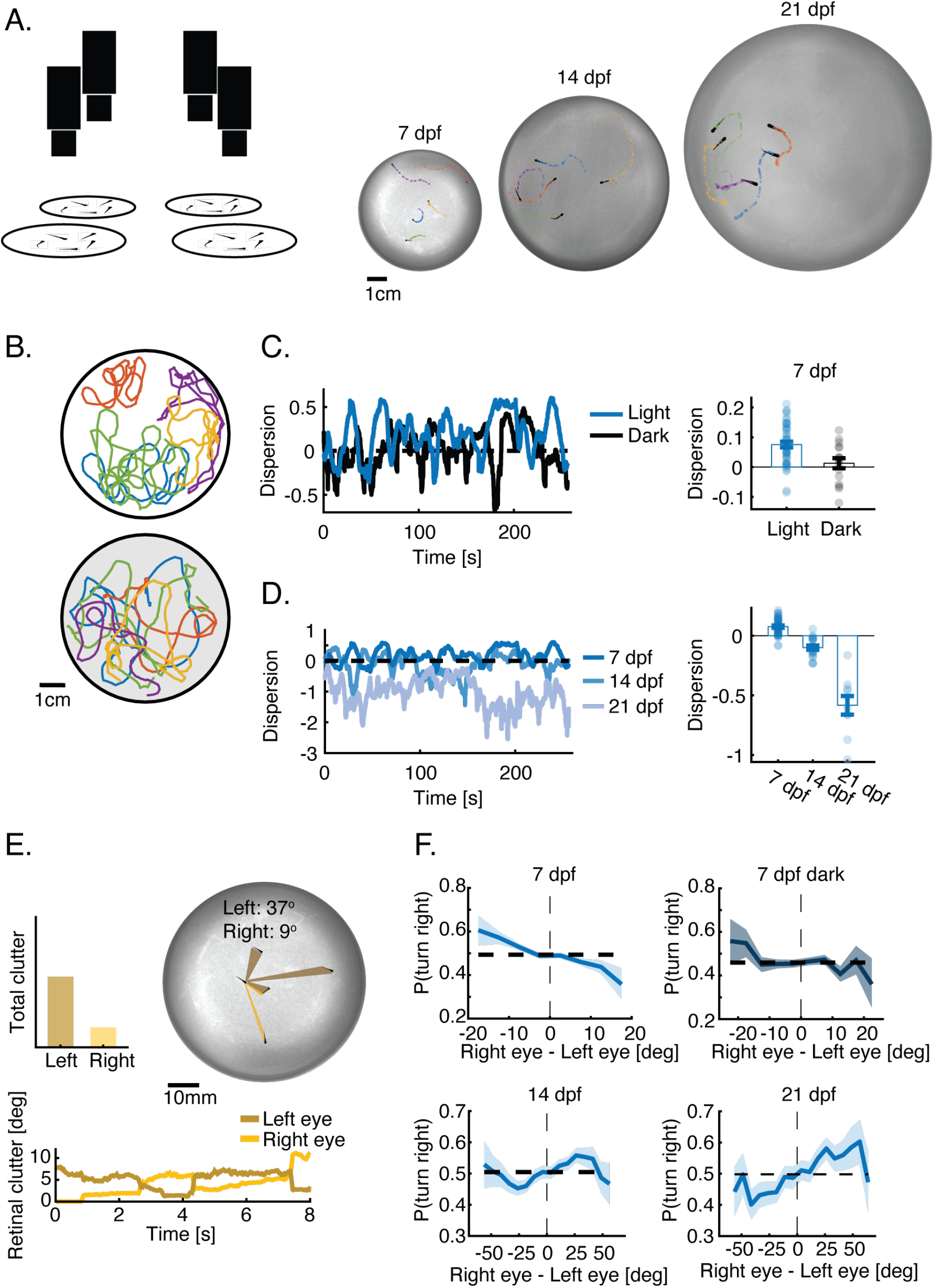
Group structure depends on visual interactions and develops with age. **A.** Left: Experimental system. Multiple cameras capture the behavior of multiple groups and individuals swimming freely in separate arenas. Right: Example images and trajectories of groups of 5 larvae at 7, 14 and 21 dpf. Different colors represent different fish in the groups. **B.** Example trajectories (recorded over 5 min) of groups of 5 larvae at age 7 dpf, swimming together in the light (top) or in total darkness (bottom). Different colors represent different fish. **C.** Left: Normalized dispersion values for one group swimming together in the light (blue) and in the dark (black)(Methods). Zero represents the average dispersion value expected when fish do not interact, and positive values represent overdispersed distributions. Right: groups of 7 dpf fish are more dispersed than is expected by chance (P<5 · 10^-9^, N_light_=48 groups, ttest) and are also more dispersed than groups swimming in total darkness (P<0.005, N_dark_ = 16 groups; ttest). Bars represent mean over groups; errorbars are SEM. **D.** Left: Example dispersion values for groups at 7, 14 and 21 dpf. Right: At 14 and 21 dpf group are significantly less dispersed (more aggregated) than chance (P<5 · 10^-4^ for both; N_14_ = 21 and N_21_ = 10; ttest), and dispersions also decreases significantly over development (P < 10^-25^, ANOVA). Bars represent mean over groups; errorbars are SEM. **E.** Top: Image showing the total angular occupancy or retinal clutter that neighbors cast on each of the eyes of a focal fish. Bottom: Example traces of the total retinal clutter for each of the eyes over 8s. **F.** Probability to turn right as a function of the difference in retinal clutter experienced by each eye (negative values - higher clutter to the left). At 7 dpf, larvae tend to turn away from the more cluttered side and do not respond to neighbors in total darkness (Top row). At 14 dpf, fish begin turning towards the more cluttered side, and this tendency increases at 21 dpf (Bottom row). Bold lines represent turning probability calculated from left/right turning events collected from all fish in 5° bins; errorbars are the 95% confidence interval of the fitted Binomial distribution to the events in each bin.

To understand how a focal fish responds to the visual information from its neighbors, we estimated the angular occupancy or retinal ‘clutter’ that neighbors projected onto the two retinae of the focal fish (57, 58)(Fig. 1E, Movie 4). We found that even a simplified global statistic of the visual input, such as the difference between total clutter experienced on each of the retinae, seemed to modulate the observed turning directions of the focal fish (Fig. 1F). Specifically, at 7 dpf, fish turned away from the more cluttered eye, and the strength of the turning response steadily increased as the difference in clutter between the retinae increased. At ages 14 and 21 dpf, on the other hand, fish turned toward the more cluttered side, and this response peaked at intermediate clutter difference values, while even larger differences in retinal activity led to a decrease of the response (Fig. 1F). No modulation of turning was observed for fish swimming in total darkness (Fig. 1F), in accordance with the lack of group structure in the dark. In addition to turning direction, we observed that the bout rate of the fish was modulated by the total integrated clutter experienced by the larvae, in which bout rate was maximal for low clutter values (Fig. S1E).

Together, these results show that visually mediated complex social interactions can be detected already at 7 dpf and that they transition from repulsive to strongly attractive by age 21 dpf. In addition, a simple global statistic representing visual occupancy on the retinae might be sufficient to explain these behaviors. Next, we use a virtual reality assay to explicitly test fish responses to retinal clutter and to infer the algorithms that allow fish to respond to complex visual scenes with multiple neighbors.

### Virtual reality reveals that young larvae specifically respond to retinal clutter

To specifically test fish responses to retinal clutter and to reveal the algorithms used to integrate information from multiple neighbors we utilized a simplified virtual reality (VR) assay, in which fish respond to projected moving objects around them, mimicking neighboring fish. We begin by focusing on 7 dpf larvae as responses in these fish are expected to be less complex than those observed in older larvae (Fig. 1). Previously, older larvae (17 dpf - 26 dpf) and adults were shown to be attracted to projected moving objects if the objects exhibit movement dynamics of real fish (31, 46). Extending these studies to 7 dpf larvae in our VR assay, we found that fish turn away from projected dots that mimic the motion of real neighbors (Fig. S2A-C, Methods), capturing both the group structure and response tendencies observed in our group swimming experiments (Fig. 1C, F).

Next, we varied the physical features, motion dynamics and number of projected objects presented to the fish, to precisely characterize their responses to these features (Fig. 2A, S3A, Methods). We generated our stimuli using a pin-hole model of the retina of the fish which transformed bottom-projected stimuli onto retinal space (Fig. 2A, S3B, Movie 5, Methods). Using this tool allowed us to independently vary specific features of the stimuli in retinal space while keeping other variables constant.

**Figure 2:**
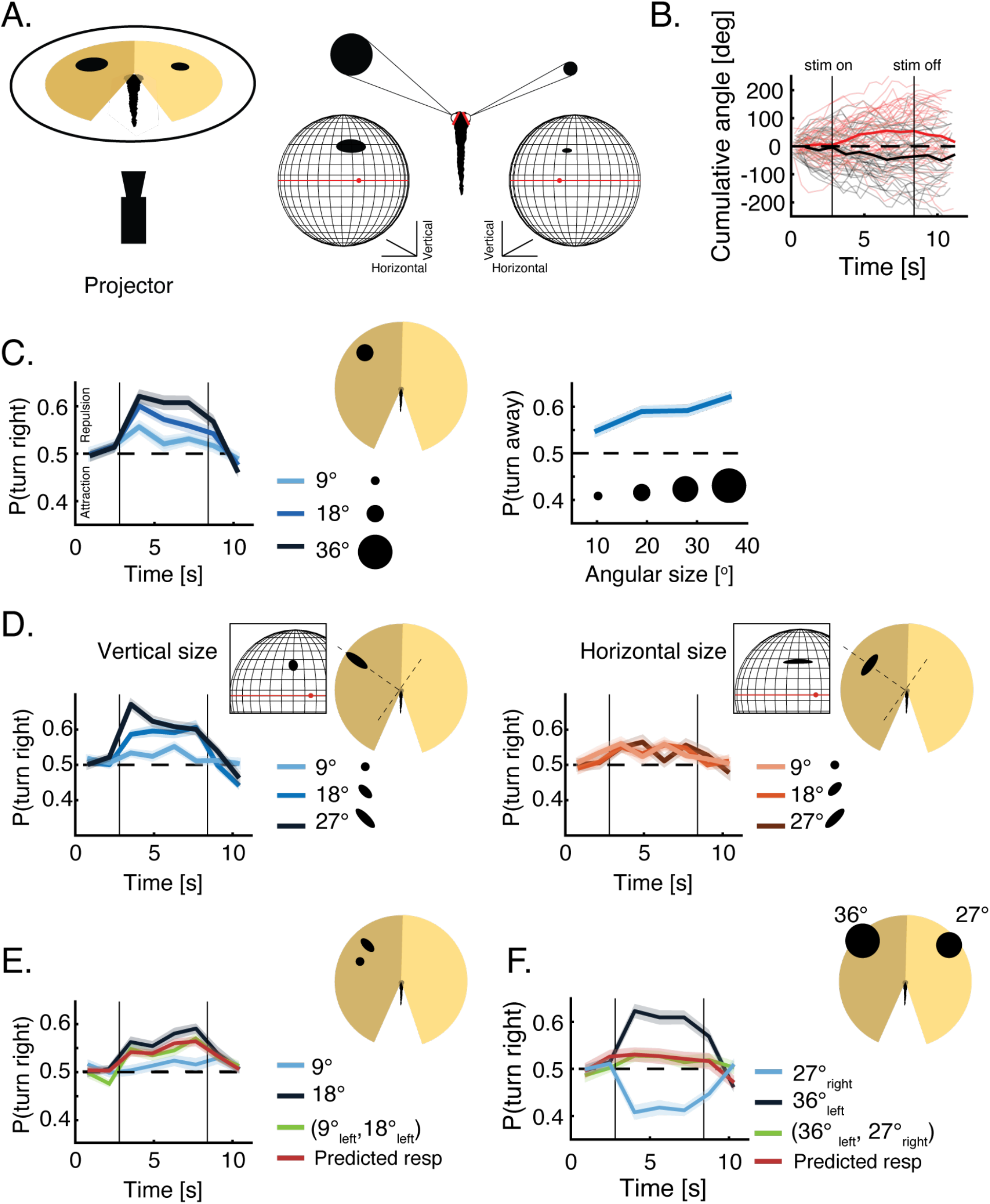
Virtual reality reveals the algorithms fish use to integrate visual social information. **A.** Left: Testing fish social interactions using closed loop projection of simplified moving objects mimicking neighbors (Methods). Right: A pinhole model of the retina is used to estimate the shape, size and position of the image of the projected object on the retina of the fish (black shapes). Both retinae are modeled as spheres, red dots are the center of the back of the retina and red lines represent the horizon line (Methods). **B.** Examples of the cumulative turning angles for one fish responding to a single dot (36°) mimicking a neighbor moving in bouts in the left visual field (red lines) and in the right one (gray lines) over 40 trials. Bold lines represent averages over trials; vertical lines represent times when stimulus is turned on and off during the trials. **C.** Left: Probability to turn right over time when a single moving dot of different sizes is presented to the left of the fish (Methods). bold lines represent the mean probability over fish, shaded areas represent SEM (N=24 fish); time is binned into 1.3s bins. Right: Average probability (over fish and trial duration) to turn away from moving dots of different sizes, presented to the left of the fish. At 7 dpf, larvae consistently turn away from the side of the projected image; shaded areas represent SEM. **D**. Left: Probability to turn right over time in response to ellipses of increasing vertical size (perpendicular to the plane of the eye), while the horizontal size remains constant at 9°. Bold lines represent averages; shaded areas are SEM (N=32 fish). Inset shows the image of the vertical ellipse on the retina. Right: Probability to turn right in response to ellipses of increasing horizontal sizes (parallel to the plane of the eye), while the vertical size remains constant at 9° (N=32 fish). Inset shows the image of the horizontal ellipse on the retina. **E.** Probability to turn right in response to two images presented together to the left visual field (green line); to each of the images presented alone (blue lines) and the prediction based on the weighted average of the responses to each stimulus presented alone (red line): *p_predicted_*(*V*^1^_*left*_, *V*^2^_*left*_) = *p*(*V*^1^_*left*_) · *w*^1^ + *p*(*V*^2^_*left*_) · *w*^2^, where V is the vertical dimension of the stimulus, and *w*^1^,*w*^2^ are weights representing the relative sizes of the stimuli such that *w^i^* = *V^i^*/Σ*V^i^*; N = 32 fish, shaded areas are SEM. **F.** Probability to turn right in response to two dots presented simultaneously to each eye of the fish (green line); to each dot presented alone (blue lines) and the prediction based on the recorded responses to each dot presented alone (red line): *p_predicted_*(*V_left_*,*V_right_*) = *P*(*V_left_*) *p*(*V_right_*) – 0.5; N = 24 fish, shaded areas are SEM.

We found that 7 dpf larvae turn away from a moving dot projected either to the left or right visual field (dots move with intermittent bouts, tangentially around the head, from ±80° to ±30°, where 0° is the heading of the fish) (Fig. 2B-C, Fig. S3A, Movie 6). Increasing the angular size of the dot, monotonically increased the probability of the fish to turn away from the stimulus, in agreement with the observed responses to retinal clutter in group swimming experiments (Fig. 2C, Fig. 1F).

Additionally, responses were similar for both stationary or moving stimuli, and repulsion tendencies completely disappeared for small objects occupying <6° on the retina (Fig. S3E).

We next tested the effects of different retinal positions by presenting stationary stimuli to different sections of the visual field, while keeping angular size constant. We found that elevation on the retina (i.e. the radial distance of the project dot) did not modulate the turning response of the fish, while the position in azimuth generated only a slight suppression at the edges of the visual field (Fig. S3C-D).

We found that fish repel away from stimuli mostly by modulating the probability of directed turns while keeping other variables such as magnitude of turns (Fig. S3F), the average path traveled in a bout and the overall bout rate constant (Fig. S3G). That lack of modulation of the average path traveled by stimulus presentation, indicates that fish responses are consistent with routine turns as opposed to large magnitude escapes (59).

These results confirm the specific role of retinal clutter in modulating the turning responses of the fish. We next test how visual information is integrated from multiple neighbors and over different dimensions of the retina to guide behavioral decisions.

### Behavioral responses to visual clutter are based on retina-wide integration and inter-eye competition

To understand how 7 dpf larvae integrate visual information over the retina we varied the physical dimensions of the projected stimuli and tested fish responses to these changes. We found that stretching the projected dot in the vertical dimension, which changes the height of the image on the retina (Fig. S3B), and increases the magnitude of vertical clutter specifically, resulted in an increased tendency to avoid the presented stimulus (Fig. 2D, left). Yet, stretching the dot horizontally, thereby changing the width of the image on the retina, and the integrated horizontal clutter, had no effect on behavior (Fig. 2D, right). The prominent role of the vertical dimension of the stimulus on the retina, was further corroborated by repeating these experiments in bowl shaped arenas, with stimuli presented to the side of the fish instead of the bottom, which allowed us to stimulate additional positions in retinal space (Fig. S4A). Importantly, we observed similar selectivity to stimuli orientation when multiple identical dots, separated from one another, were arranged vertically (i.e. same angle from the fish, at increasing radial distances) or when they were arranged horizontally (i.e. at different angles from the fish, with the same radial distance): fish increased their tendency to turn away when more dots were presented vertically, yet turned with a similar probability if one, two or three dots were presented horizontally (Fig. S4B).

To further elucidate how visual clutter is integrated from multiple objects over the retina, we presented to one eye two stimuli with different vertical sizes (and similar horizontal sizes), and found that the observed response to the combined presentation of the stimuli was an intermittent value between the two recorded responses to each stimulus presented alone (Fig. 2E). More specifically, the response to the joint presentation of the two stimuli was accurately predicted by a weighted average of the recorded responses to each stimulus presented alone, with weights equal to the relative size of the stimuli (Fig. 2E, S4C). Here again, results were similar regardless of whether the two presented stimuli were clearly separated from one another or if they were joined to create one larger stimulus (Fig. S4D). These results indicate that fish use differently the different dimensions of the retina - they integrate clutter over the vertical dimension of the retina and average the resulting values over the horizontal dimension regardless of object separation or number (see Fig. 4A below for illustration).

To understand how fish integrate visual information from both eyes, we tested fish responses when stimuli were presented simultaneously to each of the eyes (Fig. 2F, Fig. S4E and Movie 7). Here again, 7 dpf larvae tended to turn away from the side presented with the larger stimulus, yet the response tendency was attenuated compared to the case where the same stimulus was presented alone. We found that the response to two competing stimuli can be very accurately predicted by adding the two competing responses (each driving the fish in a different direction) recorded for each stimulus alone (Fig. 2F, S4E). We also note that responses to sets of stimuli that have equal angular difference between the eyes (e.g. 36° vs 27° and 27° vs 18°) were markedly different from each other, yet the response to each set could be accurately predicted using the individually recorded responses (Fig. S4E). Importantly, the attenuation caused by two competing stimuli did not seem to result in an increase in probability of forward swims that would indicate averaging of stimuli between the eyes. In fact, when two equally large stimuli were presented to both eyes, the fish were equally likely to turn away from either the right or left stimuli, which is in line with a winnertake-all strategy for binocular integration rather than averaging (39)(Fig. S4F). These results indicate that the binocular integration of the stimuli is less likely to be computed at visual sensory areas, but rather at downstream areas responsible for the behavioral responses themselves (see Fig. 5 below).

**Figure 3:**
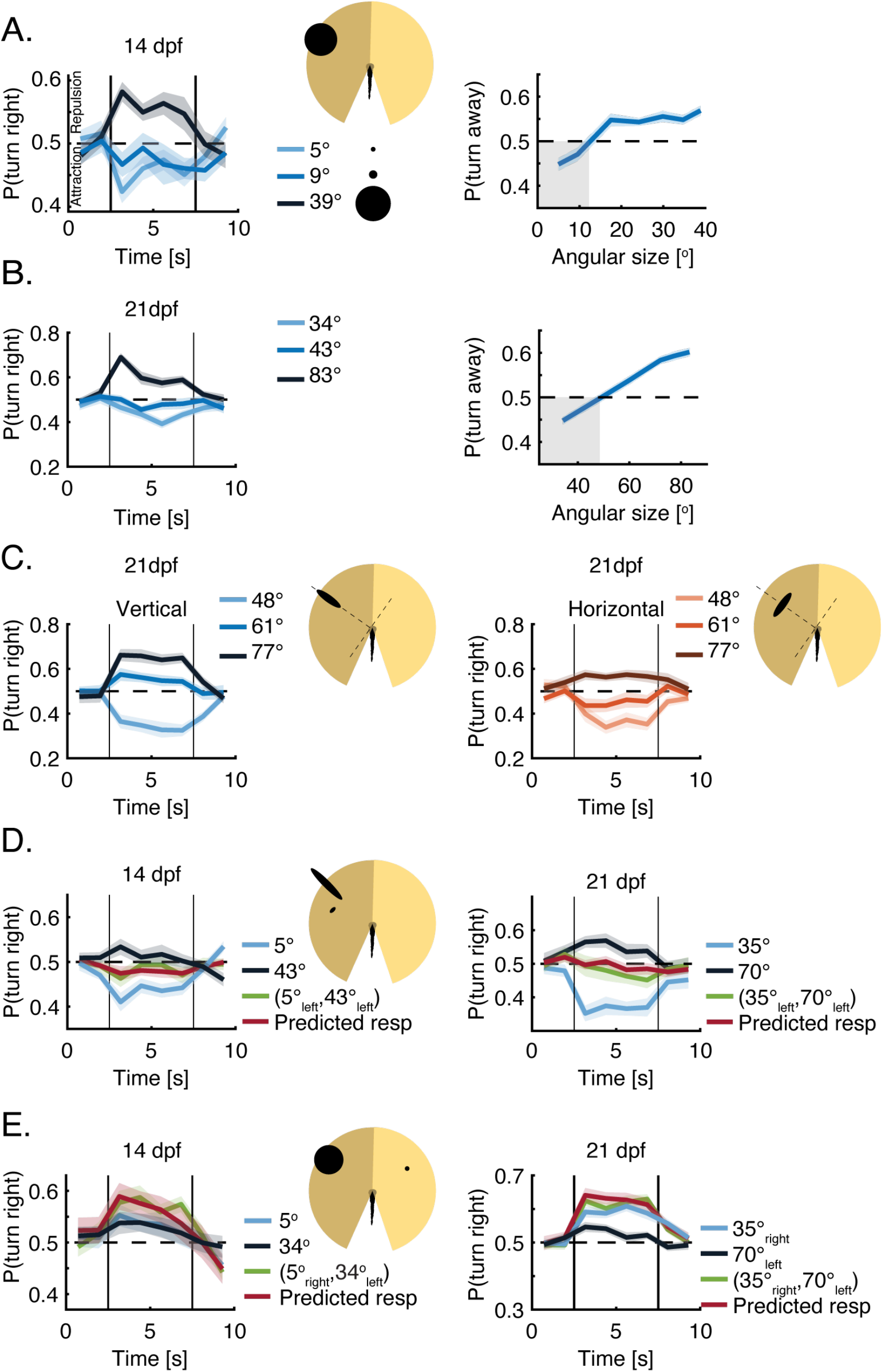
Older larvae use similar algorithms to integrate visual social information. **A.** Left: Probability to turn right over time in response to dots of different sizes presented to the left visual field. At 14 dpf, fish show both attraction to small angular sizes and repulsion from larger sizes. Bold lines represent averages, shaded areas represent SEM (N = 32 fish) and time is binned into 1.2s bins. Right: Average probability (over fish and trial duration) to turn away from dots of different sizes, presented to the left of the fish. Blue shaded area represents SEM; gray shaded area marks visual clutter values for which fish exhibit attraction **B.** Same as in A, only for 21 dpf larvae (Methods). **C.** Left: Probability of 21 dpf larvae to turn right in response to ellipses of increasing vertical size (perpendicular to the plane of the eye), while horizontal sizes remain constant at 27°. Bold lines represent averages; shaded areas are SEM (N=32 fish). Right: same but in response to ellipses of increasing horizontal sizes (parallel to the plane of the eye), while vertical sizes remain constant at 27° (N=32 fish). **D.** Left: Probability to turn right in response to two images presented together to the left visual field of 14 dpf larvae (green line); to each of the images presented alone (blue lines) and the prediction based on the average of the responses to each stimulus presented alone (red line): *p_predicted_*(*V*^1^_*left*_, *V*^2^_*left*_) = *p*(*V*^1^_*left*_) · *w*^1^ + *p*(*V*^2^_*left*_) · *w*^2^, where *w*^1^ = *w*^2^ = 0.5. Right: Same but for 21 dpf larvae. **E.** Left: Probability to turn right in response to two dots presented simultaneously to both eyes of 14 dpf larvae (green line); for each dot presented alone (blue lines) and the prediction based on the recorded responses to each dot presented alone (red line): *P_predicted_*(*V_left_*,*V_right_*) = *P*(*V_left_*) + *P*(*V_right_*) – 0.5; N = 32 fish. Right: same but for 21 dpf larvae.

**Figure 4:**
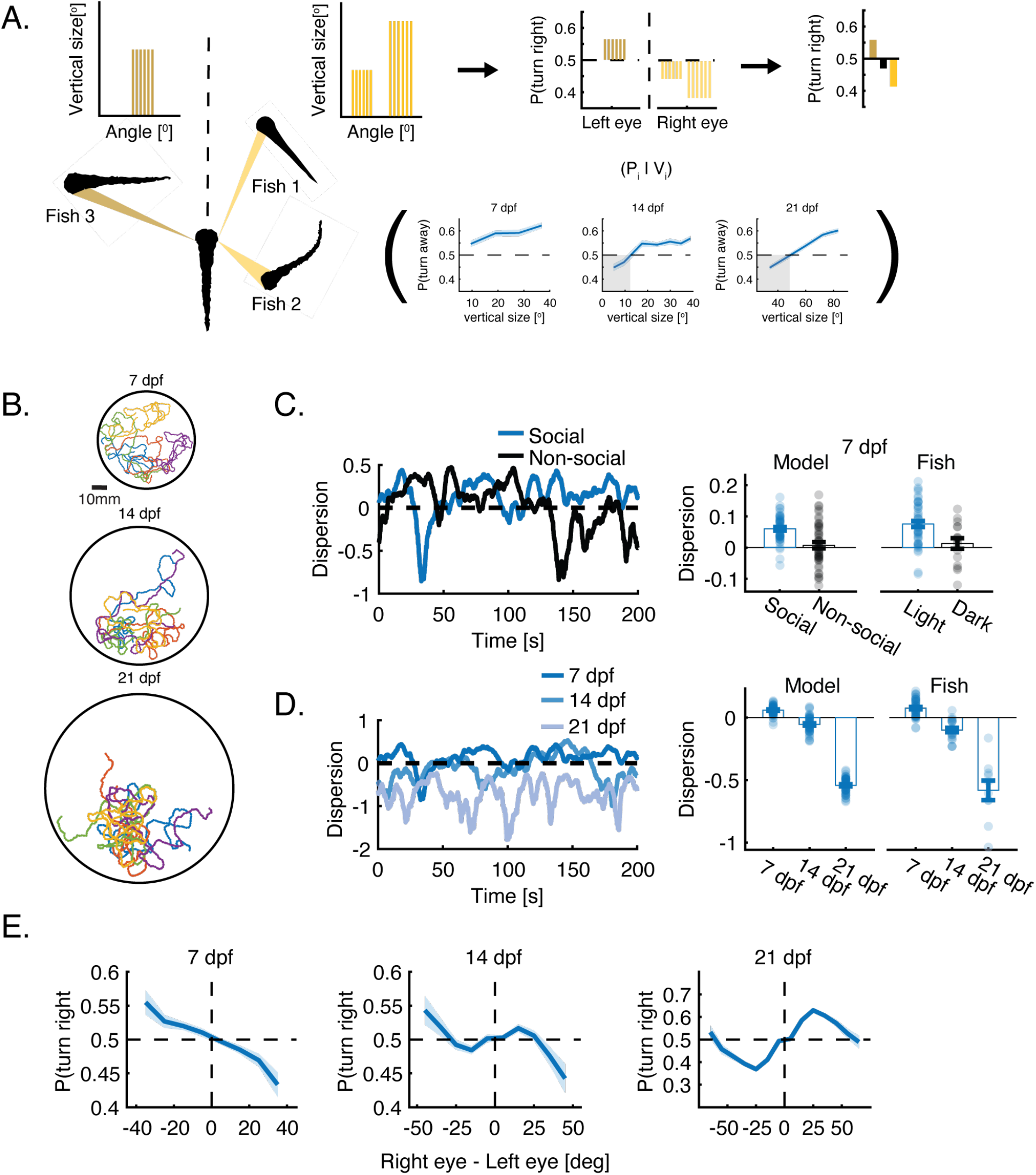
Social interactions extracted from VR capture the behavior of real groups. **A.** Models are based on vertical visual clutter casted by neighbors on the retina of the focal fish and the observed algorithms and response functions extracted from VR experiments (Eq. 1 and Methods). Left: Each neighbor is represented by its estimated vertical size at specific visual angels. Middle: The vertical sizes casted by neighbors elicit a turning bias based on the age dependent response functions observed in VR experiments (*p_i_*|*V_i_*). Right: The turning biases are then averaged on each side (yellow bars) and compared between the eyes to elicit a turing probability *p*(*turn right*) (black bar) for the next bout performed by the fish (Methods). **B.** Examples of simulated trajectories for 7, 14 and 21 dpf larvae in arena sizes that match real group experiments. **C.** Left: Normalized dispersion values for one simulated group of 5 fish at 7 dpf, with and without social interactions. Zero represents the average dispersion value expected when fish do not interact, and positive values represent overdispersed distributions. Right: Simulated groups of 5 fish at 7 dpf modelled with social interactions show higher dispersion levels than chance and are also more dispersed than simulated groups modelled without social interactions. Bars represent means; errorbars are SEM (N=50 simulations). Experimental data of fish swimming in the light and in the dark is plotted for comparison (same as Fig. 1C). **D.** Same as in C, only for simulations of different age groups. Simulated groups switch from overdispersed to clustered groups from 7 to 21 dpf. Experimental data of different age groups is plotted for comparison (same as Fig. 1D). **E.** Probability to turn right as a function of the difference in total angular occupancy experienced by each eye (negative values - higher occupancy to the left) in the simulations. (Similar to Fig. 1E-F for group experiments). Bold lines represent turning probability calculated from left/right turning events collected from all fish in 5° bins; errorbars are the 95% confidence interval of the fitted Binomial distribution to the events in each bin.

**Figure 5:**
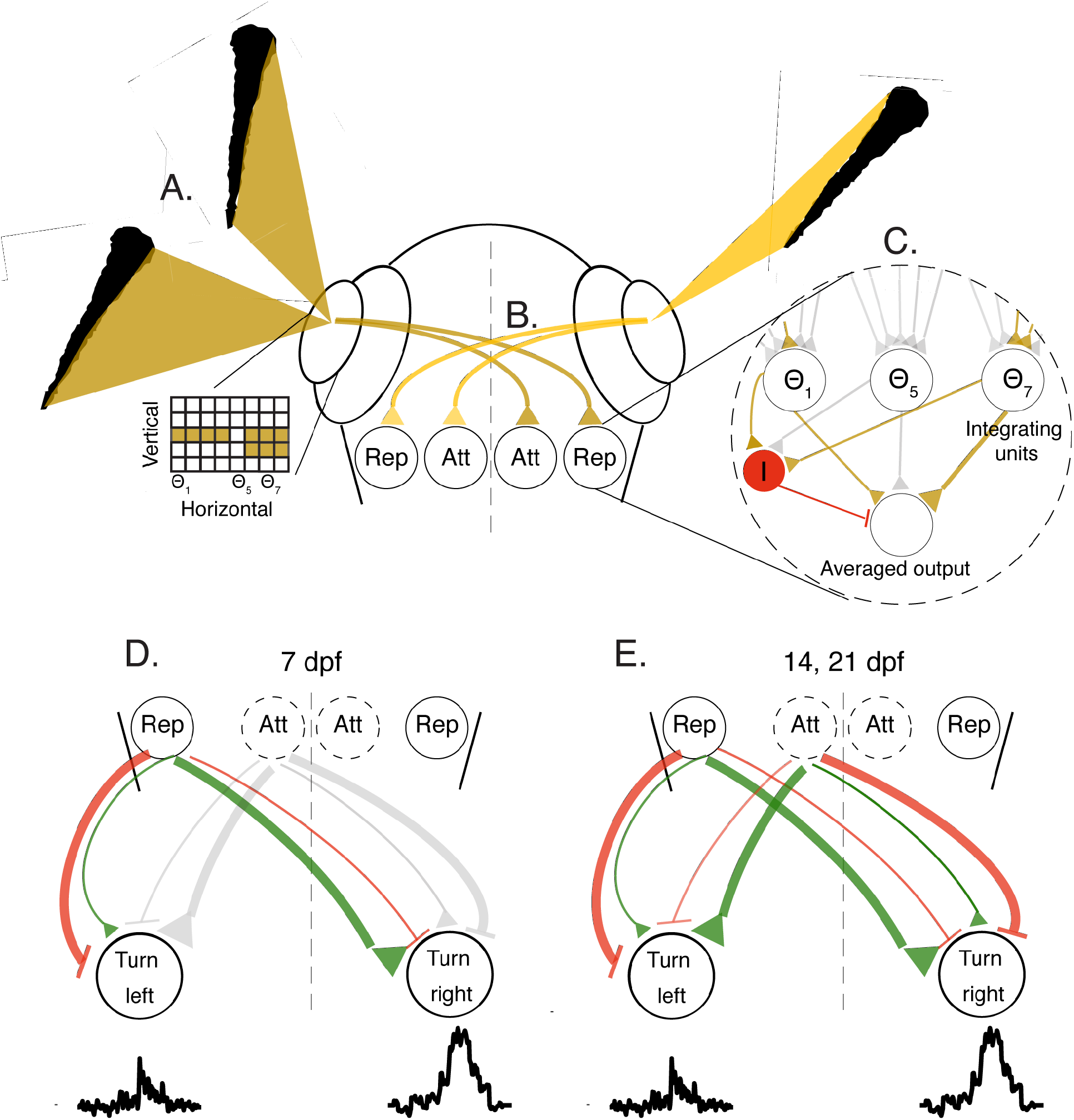
Conceptual circuit model describing visuomotor transformation underlying social interactions. **A.** Images casted by neighboring fish on the retina of the focal fish are represented as a two-dimensional grid of retinal ganglion cells mapping the vertical height of the neighbors at different viewing anglesΘ. **B.** Retinal ganglion cells selectively project to two separate populations in visual areas: the ‘Rep’ population, that ultimately elicits turning responses away from the stimulated eye, and the ‘Att’ population, that elicits turning towards the stimulated eye at older ages. **C.** The activity of all ganglion cells at a visual angle *Θ_i_* are combined by specific integrating units to represent the vertical height of the stimulus at that angle. These units relay their activity to an output population (brown triangles show activation, gray triangles represent inactive connections); we show three such example units that represent the corresponding visual angles in A. Active integrating units also stimulate an additional inhibitory population I (red) that suppresses the output population, and performs averaging of the inputs from the integrator units (see text). **D+E.** Each population in the visual areas can excite (green) and inhibit (red) motor centers on the ipsi and contralateral sides. At 7 dpf (D) responses are dominated by repulsion from the stimulated eye, while at 14 and 21 dpf (E) the balance of both the attractive and repulsive populations will determine the turning direction of the fish. Line thickness represents the strength of the excitation/inhibition, gray color represents inactive connections.

When we presented multiple stimuli together to both eyes we found, as expected, that averaging of responses to stimuli within an eye and the summation of the averaged responses between the eyes gave a very accurate prediction of the turning behavior of the fish (Fig. S4G).

Taken together, these results show that fish use different retina-wide computations to analyze visual clutter in the different dimensions of the retina: they integrate clutter in the vertical dimension, yet average over the horizontal one. Fish integrate visual information from both eyes using a winner-take-all strategy by probabilistically responding to clutter values from one of the eyes in a given response bout.

### Older larvae use similar algorithms to respond to visual clutter

We next used the VR assay to explore the way 14 and 21 dpf larvae integrate and respond to visual clutter, as fish at these ages begin turning towards their neighbors as opposed to the purely repulsive interactions at 7 dpf (Fig. 1D, F). For both these age groups, we observed the emergence of attraction to projected stimuli of small angular size, in combination with repulsion from larger stimuli (Fig. 3A-B, Methods). At 14 dpf, the transition to repulsion occurs already for very small angular sizes (< 9°), while at 21 dpf, animals remained attracted to stimuli as large as 40°, and only turned away from even larger stimuli. This is in accordance with the ontogeny of aggregation behavior in group swimming experiments in which 14 and 21 dpf larvae were increasingly attracted to their neighbors (Fig. 1D, F). Turning bias in older larvae, similar to young larvae, was mostly modulated by the probability to turn in a certain direction, while the magnitude of the turns, the bout rate and the average path traveled in a bout were only mildly affected by the size of the stimuli (Fig. S5A-B).

In line with observations at 7 dpf, the orientation of the stimulus on the retina had a marked effect on fish responses also at 14 and 21 dpf. An increase to the object’s vertical dimension (i.e. height of the image on the retina) was largely responsible for the size dependent transition from attraction to repulsion in both age groups (Fig. 3C, left). In addition, we found that unlike the 7 dpf fish, these older animals were not agnostic to changes in the horizontal dimension (i.e. width of the image on the retina)(Fig. 3C right, S5C). An increase to the width of the stimulus also contributed to its repulsive power, but to a lesser extent than an increase to its vertical dimension.

Integration of information from multiple stimuli presented together to one eye of the fish at 14 and 21 dpf, followed a similar algorithm to the one observed at 7 dpf. The responses to such joint presentation of stimuli could be accurately described by the weighted average of the recorded responses to each of the stimuli presented alone, even if the two stimuli elicited contradicting responses (attraction vs. repulsion)(Fig. 3D). Yet unlike the size dependent weighing of the stimuli at 7 dpf, equal weights to both stimuli gave the best prediction of the observed response at 14 and 21 dpf. Such equal weighing indicates that larger stimuli that elicit repulsion do not take precedence over smaller stimuli eliciting attraction, and suggest that they might involve different visuomotor pathways (see Fig. 5 below).

When presented simultaneously with stimuli to both eyes, the algorithms for binocular integration observed at 14 and 21 dpf again followed closely those seen in younger larvae. The observed response to two competing stimuli was accurately predicted by the summation of the responses to each stimulus presented alone (Fig. 3E). Interestingly, this was true also in the case where one of the stimuli evoked repulsion and the other evoked attraction, resulting in an additive effect and a higher average probability to turn in a certain direction than that of each stimulus on its own. These results suggest that while social responses become more complex as larvae mature, and involve both repulsion from larger clutter values and attraction to smaller ones, the algorithms used by 7 dpf larvae to integrate clutter over the retina are largely conserved over development.

### Modeling collective swimming behavior based on responses to visual clutter

We next tested whether social interactions based on the clutter integration algorithms extracted from VR can accurately account for group behavior in larval zebrafish. To that end, we simulated groups of 5 or 10 agents (similar to the group swimming experiments described in Figure 1) that interacted according to these rules (Fig. 4A). In total, we simulated 4 variants of the model - a nonsocial model, in which fish do not interact with one another and 3 social models (one for each age) based on the clutter integration algorithms and behavioral responses observed in VR at 7, 14 and 21 dpf (Fig. 4A, Movies 8-10). The simulated trajectories of the fish in all models were composed of discrete bouts and changes in heading direction that were based on the swimming statistics extracted from group experiments (Fig. S1D, S6A-C). In the social models, the fish biased their turning direction in each bout based on the visual clutter that the neighboring fish cast on both eyes. Specifically, each occupied horizontal visual sub-angle *θ_i_* on the retina elicits a turning bias based on its integrated vertical size *V_i_* and the age relevant turning response function learned from VR experiments - (*p_i_*|*V_i_*)(Fig. 4A). Next, these turning biases are averaged over all occupied visual angles *Θ* on each side of the fish, and finally the (signed) average responses are added, such that:

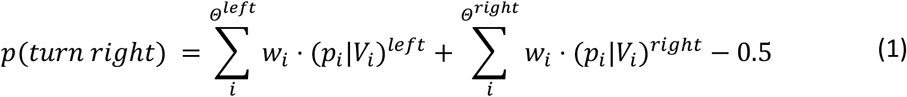

Where (*p_i_*|*V_i_*)^*left/right*^ is the turning bias elicited by the vertical size *V_i_* occupied on the retina at visual sub-angle *θ_i_* based on the age relevant response function, and *w_i_* is the relative weight assigned to that response bias (Fig. 4A). Turning direction is then set probabilistically according to *p*(*turn right*) in that bout. Thus, all models use the same clutter integration algorithms (eq. 1) yet differ in the nature of the turning bias elicited by visual clutter (*p_i_*|*V_i_*) and the relative weights *w_i_* assigned to the responses (Fig. 4A). Importantly, the models have no free parameters that are tuned to the data. Each variant of the model was simulated 50 times to account for the inherent stochasticity of the models and the results of these simulations were trajectories of moving agents in confined arenas, similar to those extracted from real groups of fish (Fig. 4B, see Methods for a full description of the models). Finally, we tested the added benefit of characterizing social interactions using VR by comparing these models to an alternative set of social models (one for each age group), that were based on the visual interactions observed in group swimming experiments (Fig. 1F). This alternative set of models were similar in all individual fish swimming properties, yet the fish in these models modulated their turning directions according to the difference in total retinal clutter between the eyes, which we estimated for all ages in the group swimming experiments (Fig. 1E-F, Methods).

### Responses to visual clutter accurately predict the behavior of real groups of fish

Simulated groups, based on the clutter integration algorithms observed in VR at 7 dpf, showed an increase in group dispersion compared to the non-social model, which exhibited dispersion values that were at chance levels (Fig. 4C). These results capture well the behavior of groups of 7 dpf larvae swimming in the light and in the dark (Fig. 4C). Simulations of 14 and 21 dpf larvae generated an age dependent decrease in dispersion (or increase in group cohesion) quantitatively similar to the pattern observed in group swimming experiments, indicating that these interactions are sufficient to explain age dependent changes in group structure (Fig. 4D). The accuracy of the models in capturing group structure also generalized well to larger groups of 10 fish swimming together (Fig. S6D). Importantly, models that were based on the algorithms extracted from the VR assay (Fig. 4A) were more accurate in predicting average aggregation of groups than models based on the visual interactions extracted directly from group swimming experiments (Fig. 1F). Average prediction errors in models based on group swimming experiments were 2.2, 1.06 and 126 times larger than those obtained by models based on VR for 7, 14 and 21 dpf fish respectively, indicating that the magnitude of the attractive responses at 21 dpf were severely underestimated in group swimming experiments (Fig. S6E). We note that simulated groups based on these minimal models did not exhibit an increase in group alignment as observed in real groups at 14 and 21 dpf, suggesting that alignment might involve additional processes not included in our models (Fig. S6F, Fig. S1B).

Our findings indicate that extracting social interactions directly from animal trajectories (as opposed to using VR) might hinder the identification of the correct interactions used by the fish (Fig. S6E). To further corroborate this finding, we attempted to extract the interaction rules used to create the simulations (Fig. 4A, eq. 1) directly from the resulting trajectories. Specifically, we repeated the calculation that was used for real fish swimming in a group (Fig. 1E-F, Methods) and estimated how the difference in total angular occupancy that simulated neighbors cast on each eye modulated the turning direction of the fish. We found that for simulations of 14 and 21 dpf fish, we could not retrieve the response functions used for the simulations, and that for 7 dpf, the inferred interactions underestimated the strengths of repulsion (Compare Fig. 4E to Fig. 4A). Yet interestingly, the (inaccurate) response functions estimated from the simulations very closely resembled those extracted from group swimming experiments (Fig. 1F). This further emphasizes that using VR can provide a more accurate description of the actual algorithms used by interacting fish.

### Constraining the underlying neurobiological circuits

The specificity of the behavioral algorithms extracted from the VR experiments allows us to make explicit predictions about the underlying neural circuits in the visuomotor processing stream. We therefore propose a conceptual circuit model, depicted in Fig. 5, that transforms visual occupancy on the retina into behavioral decisions.

This model takes the clutter elicited by neighbors on each retina as the sole input and represents it as a two-dimensional ensemble of activated retinal ganglion cells (Fig. 5A, inset). These visual inputs are relayed to downstream visual areas (e.g. optic tectum)(Fig. 5B), where a retina-wide integration of the vertical dimension is performed, thereby compressing the two dimensional grid of the retina into a one dimensional array of neurons representing the integrated values at each horizontal viewing angle (‘Repulsive’ population)(Fig. 5C). Next, the activity across this onedimensional array of cells is averaged to generate a single output value for each eye, which represents the size selective tendency of the fish to turn away from the visual clutter presented on the 2D retinal grid. Such averaging can be achieved by an additional inhibitory input to the integrating units, where the suppression is inversely proportional to the number of visual angles activated on the retina (akin to divisive normalization (60–62)(Fig. 5C).

At later stages in development (14 and 21 dpf), we propose that a second circuit module emerges, that responds maximally to a small sized vertical clutter and reduces its activity as clutter values grow (‘Attractive’ population)(63–65)(Fig. 5B). This module, by similar means, also generates a single output value for each eye that induces an inverse size selective attraction towards small clutter values. The output values from both circuit modules then excite/inhibit units in downstream areas, probably in the hindbrain, where lateralized activity is known to be responsible for controlling directed turns of the fish (43, 44)(Fig. 5D-E). At 7 dpf, these visuomotor connections are dominated by contralateral excitation and/or ipsilateral inhibition from the clutter integrating neurons (‘Repulsive’ population) to elicit competition between the two lateralized hindbrain regions, and finally a turning response away from the more cluttered eye (Fig. 5D). At 14 and 21 dpf, the additional population activated by smaller angular sizes on the retina (‘Attractive’ population) elicits an opposite response, by ipsilateral excitation and/or contralateral inhibition which results in turning towards the stimulated eye. At these older ages, the attractive and repulsive tendencies from both eyes will then compete (or add up) to elicit the observed attractive and repulsive responses of the fish.

The specific elements in this hypothesized model, e.g. units that represent integrated vertical clutter and averaged horizontal clutter in visual areas, excitation/inhibition of units in the contra/ipsilateral side in the hindbrain and even the emergence of additional modules over development can be readily tested, rejected or refined using whole-brain imaging and connectivity data from real fish (66, 67).

## Discussion

Here, we combined observations of freely swimming groups of fish with targeted manipulations of visual inputs using a VR assay, and simulations of minimal models of collective behavior to describe natural swimming behavior in groups over development and to identify the specific algorithms that govern visually based social interaction from ages 7 to 21 dpf. Our results show that larval zebrafish exhibit collective group structure already at 7 dpf and perform complex computations based on integrated retinal clutter as the input to the animal. Importantly, the basic algorithms that allow fish to integrate and respond to social visual inputs at 7 dpf were largely conserved over development, even though the repertoire of the responses to neighbors was expanded to include both attraction and repulsion at 14 and 21 dpf, as opposed to only repulsive interactions at 7 dpf. Using model simulations, we were able to show that the behavioral algorithms observed in VR experiments can very accurately describe group structure over development, which highlights the necessity of using such assays. Our findings allowed us to hypothesize the structure of the neural circuits underlying these behavioral algorithms and provide testable predictions that could be examined in future work combining our established virtual social assay with neural recordings.

Our results indicate that visual clutter is analyzed in a specific manner in which fish integrate clutter in the vertical dimension of the retina, use spatial averaging in the horizontal dimension and intereye competition based on a winner-take-all strategy to decide on the direction of their next movement. Behavioral algorithms that combine stimulus averaging and winner-take-all strategies, together with their neural substrates, were previously reported in larval zebrafish when escaping from threatening stimuli (39). The observed responses to social stimuli reported here are quantitatively and qualitatively different from escape behaviors, therefore it will be interesting to explore the similarities and differences between the brain areas and neural circuits involved in social interactions compared to those reported for escape behaviors.

Previous studies and models of collective behavior implied that animals execute complex operations such as object classification, distance measurements or object counting and that the results of these operations are available to the animals for further processing. Here we found that larval zebrafish use a much simpler strategy that does not rely on any such complex operations, namely retina-wide integration of visual clutter. These findings are in line with recent theoretical models suggesting that raw visual inputs are sufficient to elicit complex collective behaviors (10, 27, 28). Behavioral algorithms based on retina-wide integration of visual inputs are expected to fail when fish perform other behaviors such as hunting for example, which specifically requires object classification prior to any further behavioral executions. We expect such behaviors to rely on different neural circuits than the ones used for social interactions.

Interestingly, we found that visual clutter in the vertical dimension across the retina was the dominant input affecting behavior at 7 dpf, while clutter in the horizontal dimension was largely ignored. Due to a fish’s elongated shape, the horizontal extension of its projection on the retina will depend strongly on its orientation with respect to the observer. The vertical projected size, on the other hand, is less variable as it is independent of the neighbor’s orientation and only depends on its distance. We hypothesize that this is the reason why young larvae integrate only over the vertical dimension to guide their turning responses. At 14 and 21 dpf, the vertical dimension of the retinal image was still the dominant dimension eliciting repulsion, yet fish also responded to the horizontal dimension of the image comprising more complex responses. As this increase in complexity develops together with an increase in group alignment, we hypothesize that it might represent the developing tendency to detect and respond to the body orientation of neighbors as an additional input to the fish. Future experiments using VR assays as we used here, can specifically test if older larvae or juvenile fish are capable of detecting and responding to neighbors’ body orientation and motion or if they are largely agnostic to it (22, 23, 25, 26, 68).

The social responses observed in group swimming experiments and the responses we probed using the VR assay were based solely on visual input. Previous studies showed that larval zebrafish also use non-visual cues, such as mechanosensory (33, 56) and chemical stimuli (32) for social interactions. In this study we did not test how different sensory modalities operate jointly to support collective behavior. It will be interesting to test how visual information at longer distances is supported by mechanosensory sensation at shorter distances to elicit social responses (33), or how visual social information is related to chemical stimulation that represents conspecifics (32). These combinations can now be tested in future studies.

Our findings represent an important step toward elucidating the neural circuits and mechanisms at the basis of collective social behavior. First, we have detected robust computations already at 7 dpf, a critical age in which the entire nervous system of the fish is easily accessible via functional imaging techniques at single cell resolution (48–52). In addition, the basic algorithmic components we uncovered are mostly conserved during development, indicating the possibility that the underlying neural circuits are relatively matured already at 7 dpf. Second, using VR we identified the exact dimensions of the visual stimuli and the underlying algorithms that transform visual stimuli into the observed movement responses. The specificity of these algorithms allowed us to hypothesize the circuit elements involved in these computations and to make testable predictions about their structure. Performing whole brain imaging in a similar experimental assay will allow us to test, reject and refine these hypothesized circuit models, and to gain novel insight into the neural mechanisms underlying collective social behavior.

## Acknowledgments

We thank all members of the engert lab for support and advice during the project. We also thank Hanna Zwaka, Andrew Bolton, Mariela Petkova and Kristian Herrera for providing valuable feedback and suggestions in improving the manuscript and its visual content. Roy Harpaz received funding from Harvard Minds Brain and Behavior initiative. Florian Engert received funding from the National Institutes of Health (U19NS104653, R43OD024879, and 2R44OD024879), the National Science Foundation (IIS-1912293), and the Simons Foundation (SCGB 542973).

## Author Contributions

RH and FE designed research, RH, MNN and AB performed research, RH, MNN, AB and FE analyzed the data, RH and FE wrote the paper.

## Supplementary figures

**Figure S1:**
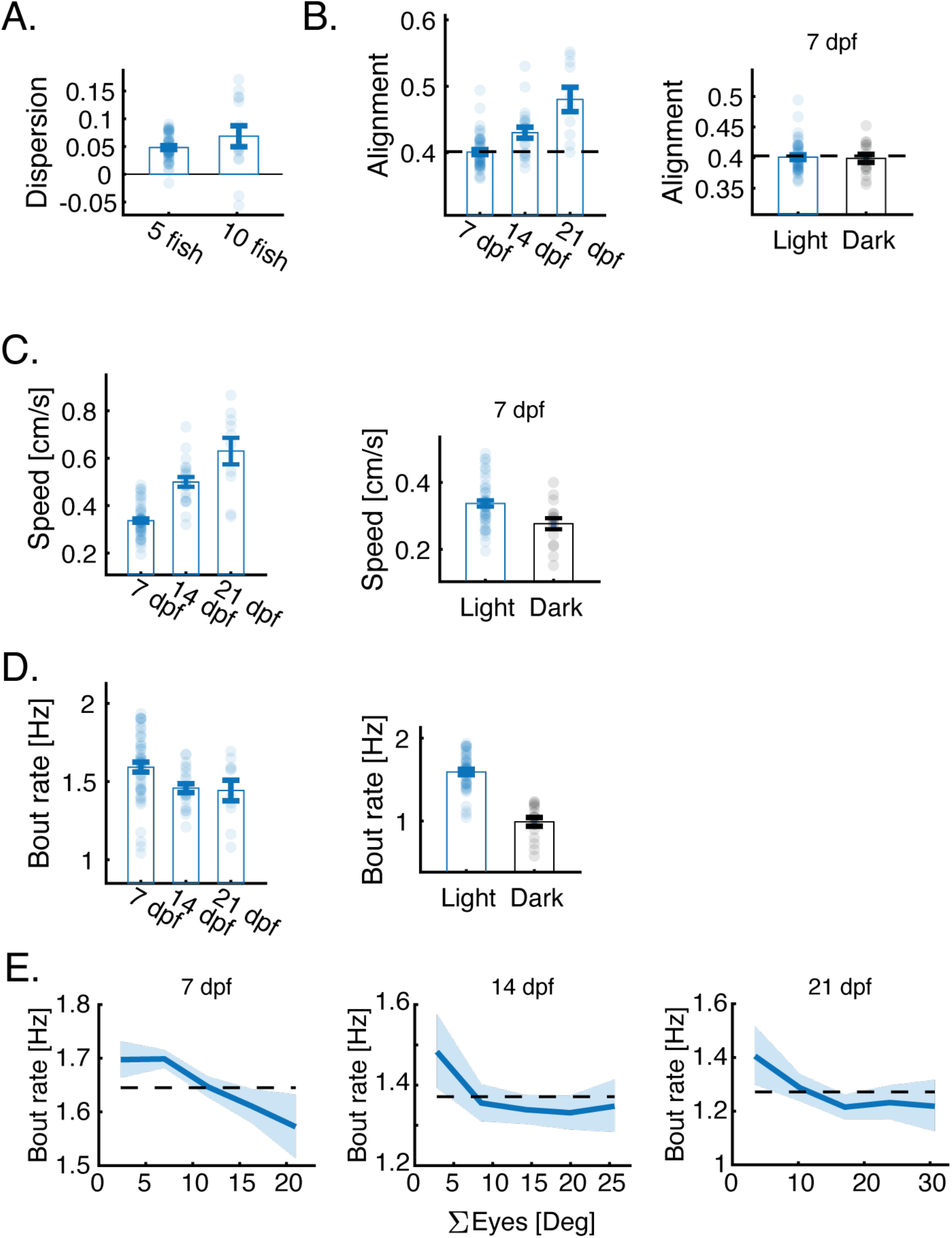
Visual social interactions develop with age. **A.** Groups of 5 and 10 fish at age 7 dpf are more dispersed than chance levels (P_5_<5 · 10^-9^, N_5_=48; P_10_ < 0.003, N_10_ = 14) **B.** Average alignment of free-swimming groups at different ages (left), and at age 7 dpf for fish swimming in the light and in the dark (right). Group alignment increases over development (P<10^-7^, ANOVA; Same groups as in Fig. 1C-D). Bars represent mean over groups and errorbars are SEM; dotted lines are group polarity values expected by chance (Methods). **C.** Same as in A, but showing the average speed of groups. Speed increases with age (P<10^-14^, ANOVA) and is decreased when 7 dpf fish swim in the dark (Right, P<0.005, ttest). **C.** Same as in B but showing the bout rate of the fish. Bout rate decreases at age 14 and 21 dpf compared to 7 dpf (Left, P<0.05, ttest), and also when 7 dpf fish swim in the dark (Right, P<10^-13^, ttest). **D.** Bout rate of the fish as a function of the total visual clutter experienced on both eyes. Fish of all ages tend to reduce their bout rate when they experience high visual occupancy. Bold lines represent the bout rate calculated over fish and groups in 5° bins; errorbars are the 95% confidence interval in each bin.

**Figure S2:**
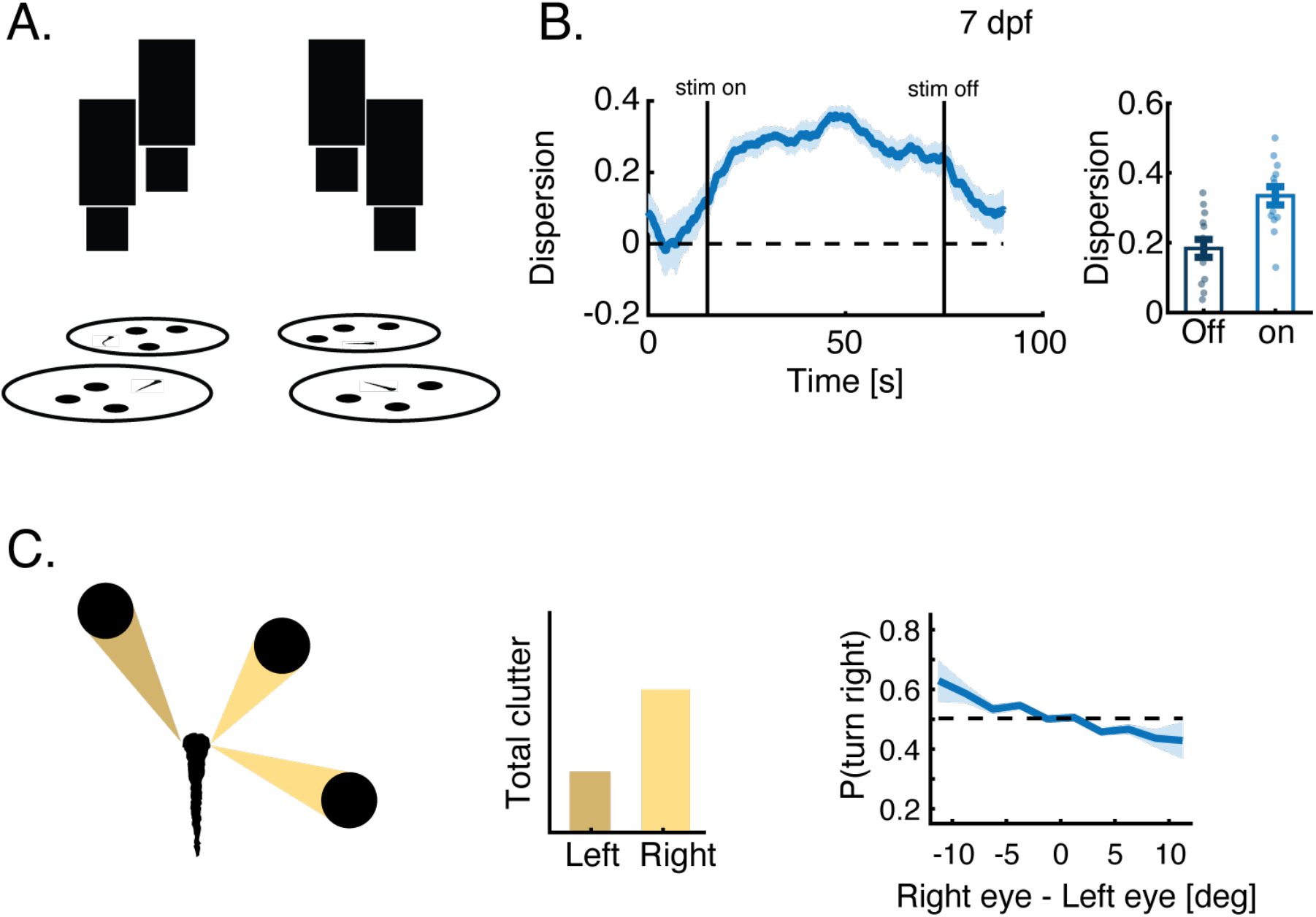
Virtual reality captures the structure and interactions of real groups at 7 dpf. **A.** Sketch of the experimental assay in which 4 fish swimming individually in different arenas are combined together via bottom projected dots mimicking the motion of real fish in separate arenas (31)(Methods). **B.** Left: Dispersion of the virtual group increases significantly when dots mimicking neighboring fish are visible to the fish (‘stim on’) compared to when they are turned off. Blue line is the average over groups and trials; shaded area is SEM. Right: Dispersion values are averaged over all times when stimulus is on vs off (P<10^-6^, N = 14 groups; ttest). **C.** Left: Sketch showing the total angular occupancy or clutter that projected dots occupy on each of the eyes of a focal fish. Right: Probability to turn right as a function of the difference in total angular occupancy experienced by each eye (negative values - higher occupancy to the left). Bold lines represent turning probability calculated from left/right turning collected from all fish in 5° bins; errorbars are the 95% confidence interval of the fitted Binomial distribution to the events in each bin. At 7 dpf, larvae tend to turn away from the more cluttered side, similar to the responses observed in group swimming experiments (Compare to Fig. 1F).

**Figure S3:**
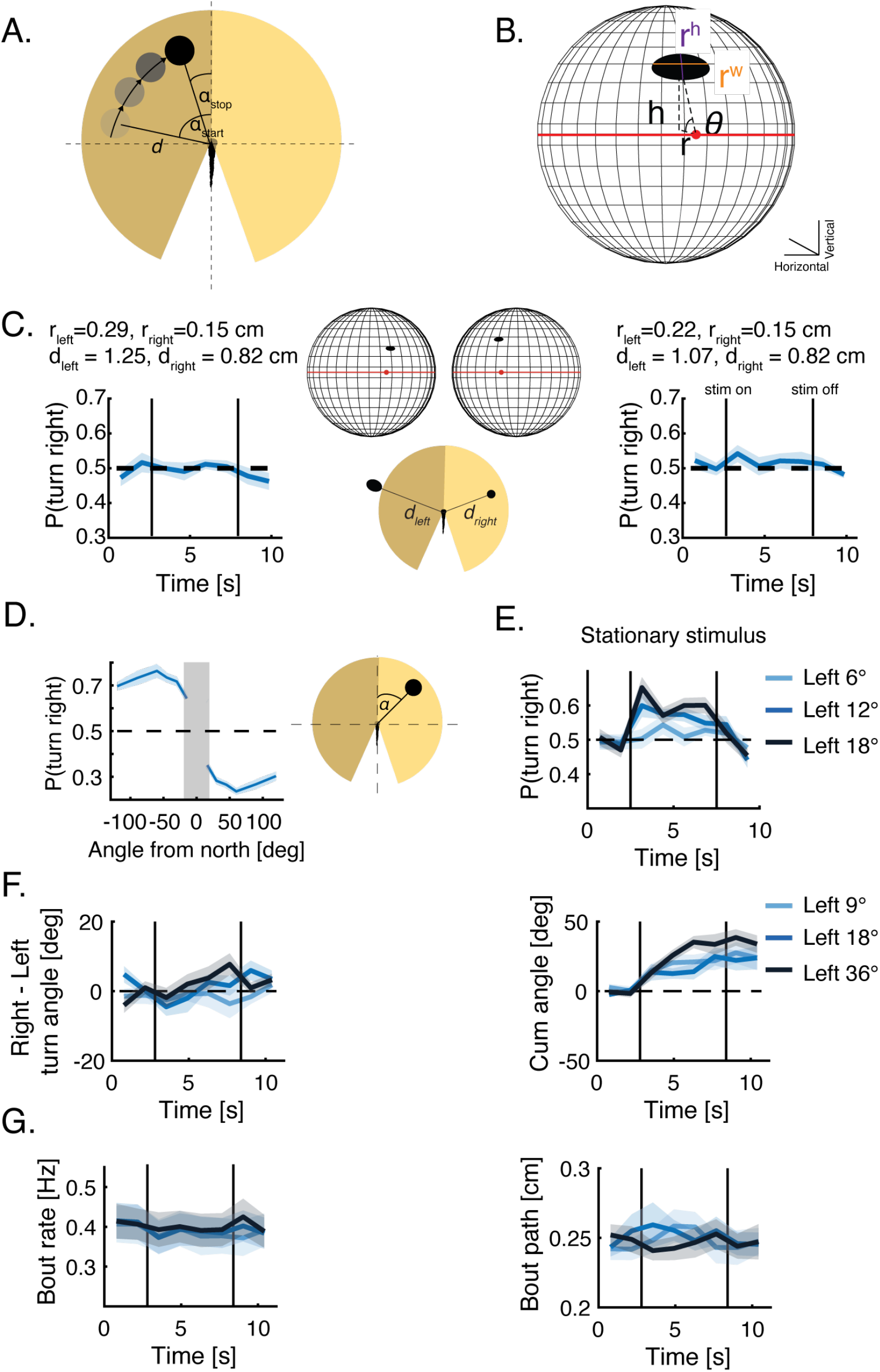
VR reveals the algorithms young larvae use to integrate visual social information. **A.** Sketch of a virtual dot moving tangentially around the fish. In every trial an image (or images) of a given size and shape appear at a starting angle *α_start_* and at distance *d* from the fish. The image moves in pancuated bouts mimicking fish motion, at a constant distance *d*, to its stopping angle *α_stop_* and disappears (Methods)(Movies 6-7). **B.** Sketch of the retina model and the calculated properties of the dot’s image on the retina (Methods). The red dot is the center of the back of the retina, and the red line represents the horizon line. *r^h^*-is the height (or vertical length) of the image, *r^w^*-is the width (or horizontal length) of the image, *h*,*r*,*θ* are the polar coordinates of the center of the image with respect to the center of the retina. **C.** Left: Probability to turn right over time when two images of different sizes *r_left_*, *r_right_*(representing the major axis of the plotted ellipse) at distances *d_left_*,*d_right_* are presented to both sides of the fish resulting in similar images on both retinae, yet at different positions. Bold lines represent average over fish, shaded areas are SEM and vertical lines represent time when stimulus is turned on and off (N=32 fish). Inset shows the projected images, sizes and distances and their retinal images (all sizes are to scale). Similar images on the retinae result in an equal likelihood to turn to left or to the right. Right: same but for different sizes and distances. **D.** Probability to turn right over time when a stationary dot is presented at different angles (and at a constant distance) with respect to the heading of the fish (0°). Positive angles are to the right of the fish. Bold line represents mean over trial duration and fish; shaded area is SEM (N=32 fish). Gray shaded area represents the expected binocular zone in the visual field that we did stimulate. **E.** Probability to turn right over time for stationary dots at different distances, presented to the left of the fish. bold lines represent the mean probability over fish, shaded areas represent SEM (N=16 fish). **F.** Difference in mean turning angle between rightward and leftward turns (Left) and cumulative turning angle (right), when dots of different sizes are presented to the left of the fish. bold lines represent the mean over fish; shaded areas represent SEM (N=24 fish). **G.** Mean bout rate (Left) and path traveled during a bout (right), when dots of different sizes are presented to the left of the fish. bold lines represent the mean over fish; shaded areas represent SEM (N=24 fish).

**Figure S4:**
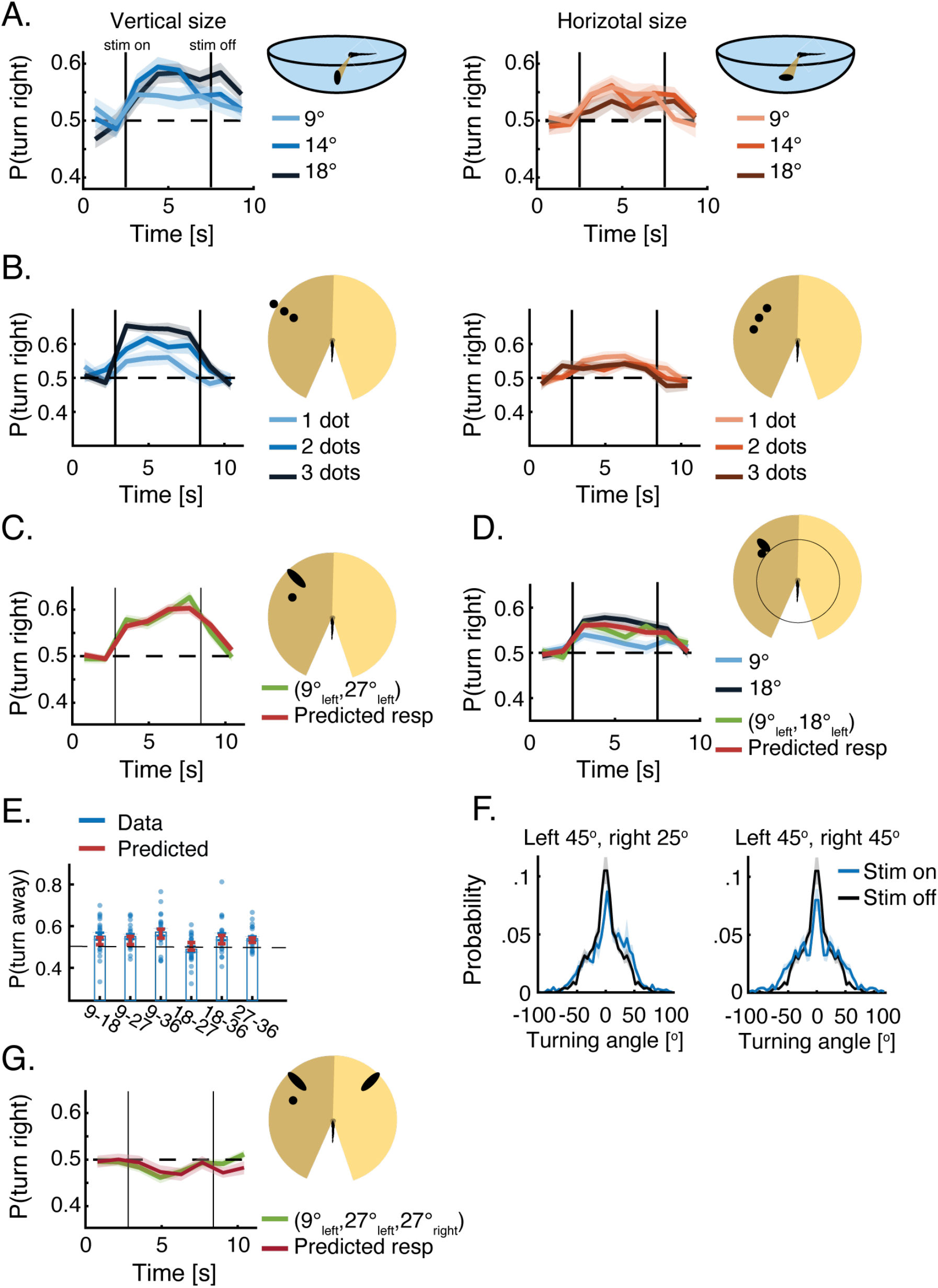
VR reveals how 7 dpf larvae integrate information from multiple neighbors. **A.** Left: Probability to turn right over time for ellipses of increasing vertical size (perpendicular to the plane of the eye), presented on the sidewall of a half dome shaped arena, while horizontal sizes remain constant at 9° (Methods). Bold lines represent averages; shaded areas are SEM (N=32 fish). Right: Probability to turn right for ellipses of increasing horizontal sizes (parallel to the plane of the eye) presented on the sidewall of a half dome shaped arena, while vertical sizes remain constant at 9° (N=32 fish). Results are similar to those observed for ellipses presented on the bottom of the tank (Fig. 2D). **B.** Left: Probability to turn right when one, two or three dots of similar size (9°) are presented to the left visual field. Dots are positioned at different angles but with a similar radial distance from the fish. Bold lines represent averages; shaded areas are SEM (N = 32). Right: Probability to turn right when one, two or three dots of similar size (9°) are presented to the left visual field. Dots are positioned at the same angle from the fish, but with different radial distances (N = 32). Probability to turn right increases with the increase in vertical clutter but not with the increase in horizontal clutter (same as Fig. 2D for ellipses of different sizes). **C.** Similar to Fig. 2E but with a different combination of stimuli (N = 32 fish). **D.** Similar to Fig. 2E, but with no space between the two images presented to the left of the fish. Larvae respond in a similar manner when the two stimuli are separated from one another and when they are not (N=32 fish). **E.** Mean probability to turn right when two dots of different sizes are presented to both eyes of the fish (larger dot is always presented to the left visual field). Bars represent mean over groups (blue dots); errorbars represent SEM. Overlaid are the mean and the SEM of the predicted responses of the groups based on the observed response to each of the stimuli presented alone (red bars)(see Fig. 2F). **F.** Distribution of fish turning angles when two stimuli of different sizes (Left) and of similar sizes (Right) are presented on both sides of the fish (blue). For comparison, the distribution of fish turning angles without any stimuli is also shown (black). Turning angles for each fish (N=8 fish) are grouped into 5° bins over all trials (Methods). Bold lines represent mean over fish; shaded areas are SEM. The distributions of turning angles indicate that fish do not average stimuli sizes, but rather probabilistically respond to one of the two presented stimuli in each bout. **G.** Probability to turn right when two images are presented to the left eye and a single image is presented to the right one (green line)(N = 32 fish). The response to the joint presentation of the stimuli is accurately predicted by averaging the responses obtained when the images are presented alone within an eye and taking the difference of the averaged responses between the eyes (red line). *p_predicted_*(*V*^1^_*left*_,*V*^2^_*left*_,*V_right_*) = *p*(*V*^1^_*left*_) · *w*^1^ + *p*(*V*^2^_*left*_) · *w*^2^ + *p*(*V_right_*) – 0.5 where V is the vertical dimension of the stimulus, and *w*^1^, *w*^2^ are weights representing the relative sizes of the stimuli such that *w^i^* = *V^i^*/Σ*V^i^*; N = 32 fish, shaded areas represent SEM.

**Figure S5:**
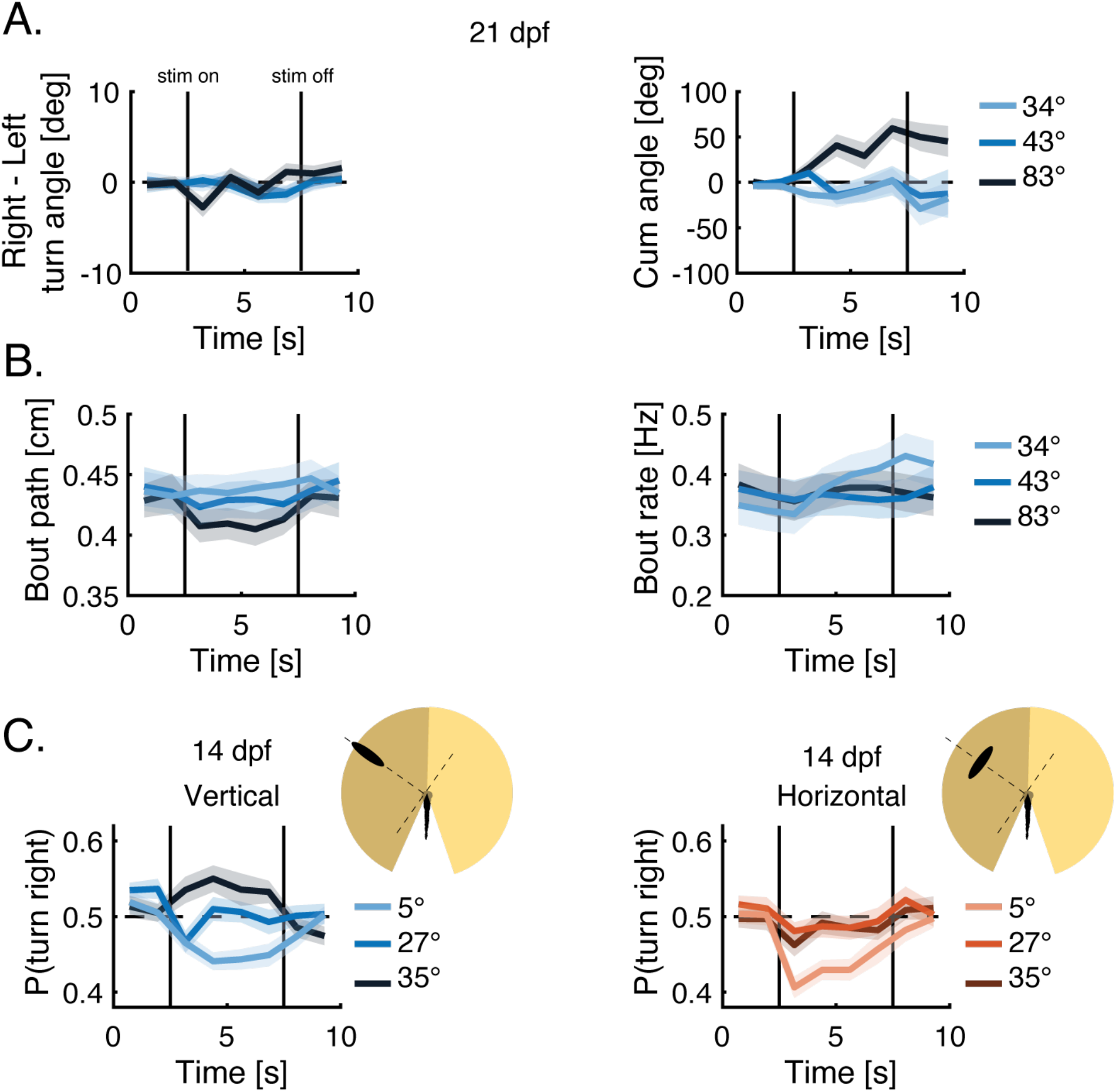
VR reveals that older larvae use similar algorithms to those seen in young larvae. **A.** Difference in mean turning angle between rightward and leftward turns (left) and cumulative turning angles (right), when images of different angular sizes are presented to the left of 21 dpf fish (Methods). bold lines represent the mean over fish, shaded areas represent SEM and vertical lines represent times when stimuli are turned on and off (N=32 fish). **B.** Mean bout rate (left) and path traveled during a bout (right), when images of different angular sizes are presented to the left of 21 dpf fish. bold lines represent the mean over fish, shaded areas represent SEM (N=32 fish). **C.** Left: Probability to turn right in response to ellipses of increasing vertical size (perpendicular to the plane of the eye), while horizontal sizes remain constant at 9°. Bold lines represent averages; shaded areas are SEM (N=32 fish). Right: Probability to turn right in response to ellipses of increasing horizontal sizes (parallel to the plane of the eye), while vertical sizes remain constant at 9° (N=32 fish, age 14 dpf).

**Figure S6:**
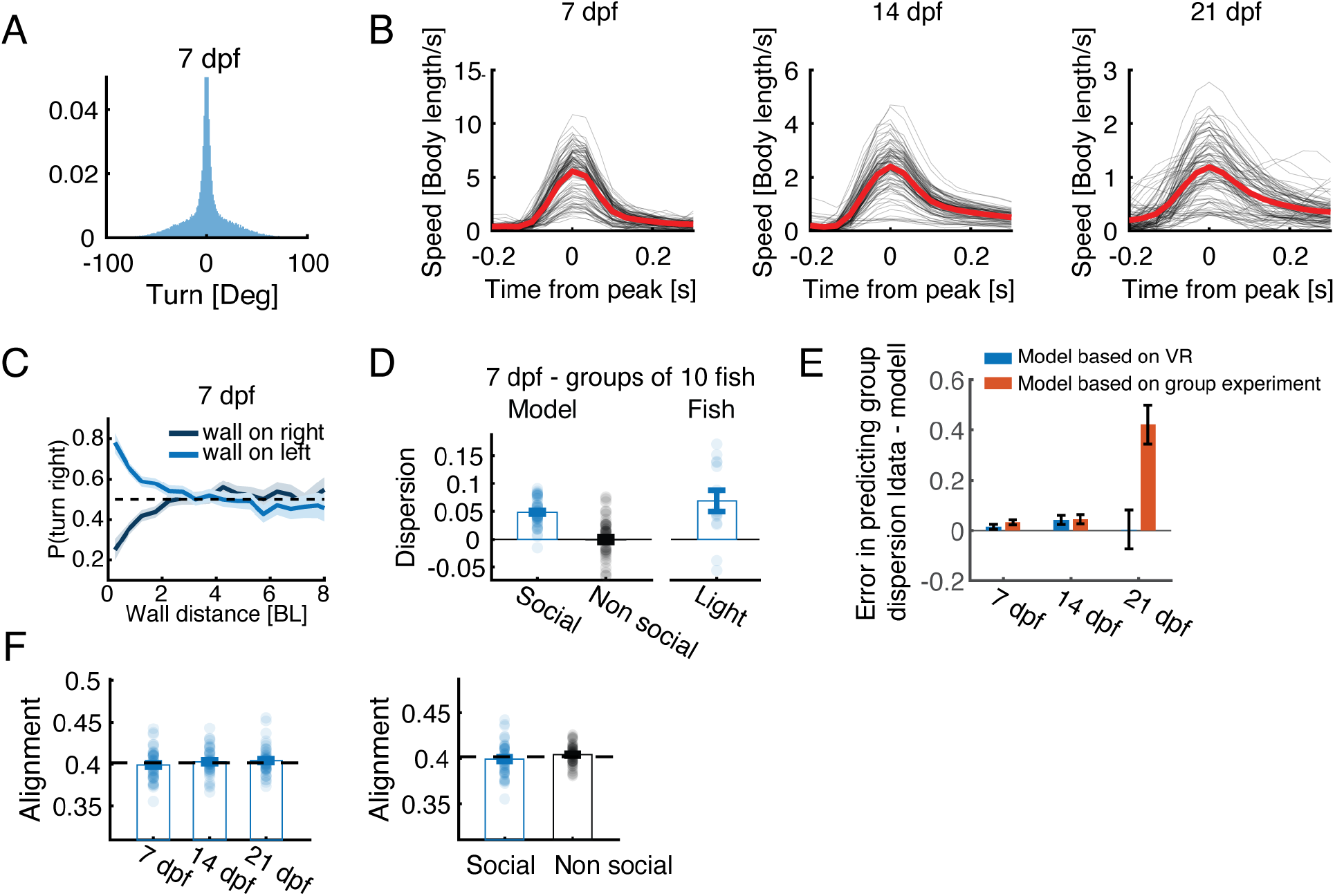
Modeling collective behavior based on responses to visual clutter from VR. **A.** Distribution of turning angles estimated from all experiments of 7 dpf fish swimming in the light (N = 48 groups of 5 fish). **B.** Speed profiles of real fish at different ages (black) and the average observed profile (red)(Averages are based on 100 bouts extracted from the different age groups). Speed profiles in the simulations were taken to approximate the average path travelled in a bout for the different ages (Methods). **C.** Probability to turn right as a function of the distance and direction (left/right) to the closest wall. Bold line is the probability estimated from right/left turns collected from all fish in 1 body length bins; shaded area is the 95% confidence interval of the fitted Binomial distribution to each bin. Fish consistently turn away when swimming close (< 3 body lengths) to a wall. **D.** Simulated groups of 7 dpf fish (10 fish in a group) with social interactions are more dispersed than is expected by chance and are also more dispersed than simulated groups without social interactions. Bars represent means; errorbars are SEM (N=50 simulations). Experimental data of groups of 10 fish swimming in the light is plotted for comparison (N = 14). **E.** Mean error in predicting the average dispersion of real groups for models that are based on social interactions extracted from VR assay (blue, Fig. 4A) and for models that are based on the interactions inferred directly from group swimming data (red, Fig. 1F). Errorbars represent SEM based on the sample sizes of group experiments (Fig. 1D). **F.** Mean polarity values for simulated groups of fish with social interactions for different age groups (left) and for simulated 7 dpf fish with and without social interactions (right). Alignment values in all simulations are at chance levels.

## Movies

**Movie 1-3: Free swimming behavior of larvae in a group.** Example groups of 5 larvae swimming together at ages 7 dpf (Movie 1), 14 dpf (Movie 2) and 21 dpf (Movie 3). Colors represent individual fish; movies are shown at x2 real speed.

**Movie 4: Estimating retinal clutter using ray casting.** Example showing the estimated retinal clutter that each neighbor occupies on the eye of a 7 dpf focal fish (right), together with total clutter experienced on each eye (left). Yellow cones represent clutter that neighbors occupy on the right eye and brown cones on the left eye. Movie is shown at x5 real speed.

**Movie 5: Estimating projected retinal images using a pinhole model of the retina.** Examples showing two projected dots (left), sizes 36° and 9°, at 4 distinct angles around the head of the fish from ±80 to ±30 (the eyes of the fish and the planes of the retinea are also shown) and their corresponding retinal images (Right). Red dots represent the centers at the back of the retinae.

**Movie 6: Fish responding to a closed-loop monocular stimulus.** Example trial of a fish responding to a dot of size 36° moving in intermittent bouts tangentially to its right (from 80° to 30°, where 0 is the fish’s heading direction). Trials begin with 2.8s without a stimulus, followed by 5.6s of stimulus presentation and end with an additional 2.8s without a stimulus. Stimulus is presented to the fish only at times when the fish is stationary.

**Movie 7: Fish responding to closed-loop binocular stimuli.** Example trial of a fish responding to two dots moving in intermittent bouts tangentially around its head to the left (size 9°) and to its right (size 36°)(from ±80° to ±30°, where 0 is the fish’s heading direction). Trials begin with 2.8s without stimuli, followed by 5.6s of stimuli presentation and end with an additional 2.8s without stimuli. Stimuli are presented to the fish only at times when the fish is stationary.

**Movie 8-10: Simulated groups based on algorithms extracted from VR.** Examples of 5 simulated fish swimming in groups at ages 7 dpf (Movie 8), 14 dpf (Movie 9) and 21 dpf (Movie 10). Simulated fish interact with one another based on the algorithms extracted from VR experiments (Fig. 4A and Methods). Colors represent individual fish

## Methods

### Fish husbandry

All larvae used in the experiments were obtained by crossing adult AB zebrafish. Larvae were raised in low densities of approximately 40-50 fish in large petri dishes (D=12cm). Dishes were filled with filtered fish facility water and were kept at 28°c, on a 14-10h light dark cycle. From age 5 dpf, fish were fed paramecia once a day. On day 7, fish that were not tested in behavioral experiments, were returned to the fish facility where they were raised in 2L tanks filled with 1.5” nursery water (2.5ppt), with ~15 fish in each tank and no water flow. On days 10-12 water flow was turned on and fish were fed artemia 3 times a day until they were tested at 14 or 21 dpf. All experiments followed institution IACUC protocols as determined by the Harvard University Faculty of Arts and Sciences standing committee on the use of animals in research and teaching

### Free swimming experiments

Fish were transferred from their holding tanks to custom-designed experimental arenas of sizes d=6.5,9.2,12.6 cm, depending on the age of the fish (7, 14 and 21 dpf respectively) filled with filtered fish facility water up to a height of ~0.8 cm. Experimental arenas were made from 1/16” clear PETG plastic and had a flat bottom and curved walls (half a sphere of radius 0.5 cm) to encourage fish to swim away from the walls. Arenas were sandblasted to prevent reflections. Every experimental arena was filmed using an overhead camera (*Grasshopper3-NIR*, *FLIR System*, *Zoom 7000*, *18-108mm lens*, *Navitar*) and a long pass filter (R72, Hoya). All experimental dishes were lit from below using 6 infrared LED panels (940 nm panels, Cop Security) and by indirect light coming from 4 32W fluorescent lights. Every 4 cameras were connected to a single recording computer that recorded 4MP images at 39fps per camera. To prevent overload of the RAM we performed online segmentation of the recorded images and saved only a binary image from the camera stream. The segmented images were then analyzed offline to extract continuous tracks of the fish using the tracking algorithm described in (29). All acquisition and online segmentation were performed using costume designed software written in Matlab. Groups were eliminated from subsequent analysis in the case that one or more of the fish were immobile for more than 25% of the experiment. All and all 35%, 22% and 33% of groups ages 7, 14 and 21 dpf were eliminated from the analysis due to immobility of the fish. Choosing a more stringent, or a less stringent criteria for elimination did not change the qualitative nature of the results.

### Individual and group properties of free-swimming fish

The position of each fish *i* at time *t* is denoted as 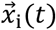. The velocity of each fish *i* is given by 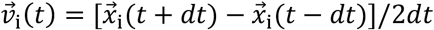, where dt is 1 frame or 0.025s. The speed of the fish is then 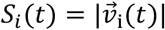, and the direction of motion is 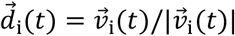.

For the group, we calculate a normalized measure of group dispersion: *Dispersion* = *log*(*NN*_1_/*NN*_1_^*shuffled*^) where *NN*_1_ is the average nearest neighbor distance and *NN*_1_^*shuffied*^ was calculated from randomized groups created by shuffling fish identities such that all fish in a given randomized group were chosen from different real groups. Positive dispersion values mean that real groups are more dispersed than shuffled controls and 0 means equality. Group alignment was defined as 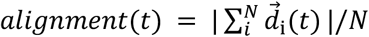, where N is the number of fish in the group. Chance levels were calculated from randomized groups (see above), and alignment values are bounded between 0 (all fish are facing in different directions) and 1 (fish are completely aligned).

### Estimating retinal clutter using ray casting

To estimate the retinal clutter or visual angle that each neighbor in the group casts on the eye of focal fish *i*, we used a modified ray casting algorithm (53, 57). Specifically, we casted 1000 rays from each eye of the focal fish spanning 165° from the direction of motion towards the back of the fish, leaving a total of 30° of blind angle behind the fish. This amounts to an angular resolution of ~0.165° per line. We then detected all pixel values representing fish in the paths of the rays and calculated the visual angle occupied by each fish and the total occupied visual angle experienced by each eye (Fig. 1E).

### Segmenting fish trajectories

Trajectory segmentation into discrete bouts or decision events was done by detecting local minima points in the speed profile of the fish (29). A bout was defined as the motion between two consecutive local minima. Individual events were then characterized by the duration of the event, the total path traveled and the change in angle, or turning response between the start and the end of the event.

### Turning in response to the arena walls

To estimate how the walls of the arena affect the turning behavior of the fish we calculated the probability of the fish to turn in a certain direction for a given distance (*D_wall_*) and direction (left/right) of the closest wall: *P*(*turn right*\*D_wall_*)^*left/right*^ (Fig. S6C). Distance to the wall was grouped into 1 body lengths bins, and turning probability was calculated from all events, pooled together from all fish, in a given bin. Errorbars represent the 95% confidence interval of the fitted Binomial distributions to the data in each bin. Responses to the wall seem to decay to chance levels at distances > 3 body length.

### Turning in response to the difference in retinal clutter between the eyes

We estimated how the difference in total retinal clutter between the eyes *Δvisual clutter* (see above) affects the binary turning direction (either left or right) of fish swimming in a group (Fig. 1F). Specifically, we estimated *P*(*turn right*|*Δvisual clutter*) which is the probability to turn in certain direction for 5° bins of *Δvisual clutter* and estimated the 95% confidence interval from the fitted Binomial distribution to the data in each bin. We discarded all turning events at distance < 3 body length from the wall, as not to confound wall avoidance with neighbor responses. Data is pooled over all fish.

### Effects of total retinal clutter on bout rate in groups of fish

We estimated how the total retinal clutter in both eyes *Σvisual clutter* affects the probability of the fish to perform a bout or a movement decision (Fig. S1D). Specifically, we estimated *P*(*bout*\*Σvisual clutter*) which is the probability of a bout for 5° bins of *Σvisual clutter* and estimated the 95% confidence interval from the fitted Binomial distribution to the data in each bin. We discarded all bouts at distance < 3 body length from the wall, as not to confound wall avoidance with neighbor responses. Data is pooled over all fish.

### Virtual reality assay

We combined the experimental system that was used to track groups of fish together with bottom projection of visual stimuli in closed loop as our virtual reality assay (Fig. 2A). All experiments were done using experimental arenas with a diameter of 9.2cm with a single fish in each arena interacting with images projected directly onto the sandblasted flat bottom of the arena (Fig. S3A). All fish tracking and posture analysis were done using custom software written in Python 3.7 and OpenCV 4.1 as described extensively in (43). Briefly, movie images acquired at 90Hz were background subtracted online to obtain an image of the swimming fish, and body orientation was estimated using second-order image moments. We used the specific position and body orientation of the fish to present moving images that are locked to the position and heading direction of the fish (Movies 6-7). We defined swim bouts using a rolling variance of fish orientation (50 ms sliding window) with a constant threshold. Visual stimuli were presented only when fish were stationary and were turned off during a bout.

### Visual stimuli

Images were presented on one or both sides of the fish. Stationary images appeared at a constant angular position (±50° from the heading of the fish) and radial distance (0.825 cm to the closest edge of the presented image) with respect to the fish and stayed on while the fish was not performing a bout and until the end of the trial. Different trials were separated by an inter-stimulus interval (ISI) equal in length to the time of stimulus presentation (5s). The temporal order of different stimuli was randomly shuffled during an experimental session.

Moving images appeared at a constant radial distance (0.825 cm to the closest edge of the image) in the periphery of the visual field (±80°with respect to the fish’s direction of motion 0°) and moved towards the center of the visual field (±30°) in bouts mimicking fish natural motion (Movies 6-7).

For 21 dpf larvae, due to the larger range of angular sizes needed to probe the behavior of the fish, visual occupancy was modulated by positioning a constant size image (0.8 cm in diameter) at different radial distances (instead of changing the size of the image presented at a constant distance, as was done for 7 and 14 dpf fish). When more than one image was presented to the same visual field of the fish, unless otherwise stated, the images were separated by empty space equal to the width of the presented images.

In all experiments, every stimulus or stimulus combination presented to the fish always had its mirror image stimuli presented on a separate trial. In all analyses throughout the manuscript these mirror image trials are flipped and combined together.

### Virtual interactions in a bowl-shaped arena

We projected images (on one or both sides of the fish) onto a half dome shaped arena (R = 3.6cm) made from commercially available light diffusers (*Profoto).* Domes were filled with water to the top, and projected images were centered at the midlevel of the dome, i.e. ~1.8 cm from the bottom. Projected images were corrected to account for the curvature of the dome and to eliminate distortions. We used stationary stimuli situated at ±60 degrees from the fish’s heading direction. We changed the size of the projected image depending on the distance of the fish to the walls, such that the estimated angular sizes of the vertical and horizontal axis were constant. The maximal angular size we used wasl8°degrees to avoid images becoming too large when the fish is far from the wall. We did not present images when the fish’s distance from the center of the arena was larger than the distance of the middle of the projected image from the center of the area. In these cases, the fish was too close to the walls, and we could not estimate the size of the projected image.

### Measuring virtual interactions between fish swimming in separate tanks

We tracked the positions of 4 individual fish, each swimming in a separate identical arena (D = 9.2 cm). We then projected 3 moving dots (D = 0.3 cm) in each of the separate arenas, that exactly mimicked the position and velocity of the 3 fish swimming in the other arenas (Fig. S2A)(31). Every experiment consisted of 60 trials, and each trial consisted of 60 seconds where the dots were visible to the fish (‘on’) and 30 seconds where the dots were not visible (‘off’). We then collected the tracked position of the 4 real fish from the separate arenas and analyzed them as a single group (Fig. S2).

### Retina model

To estimate the shape, size and position of the projected object’s image on the retina of the fish, we used a pinhole model of the retina:

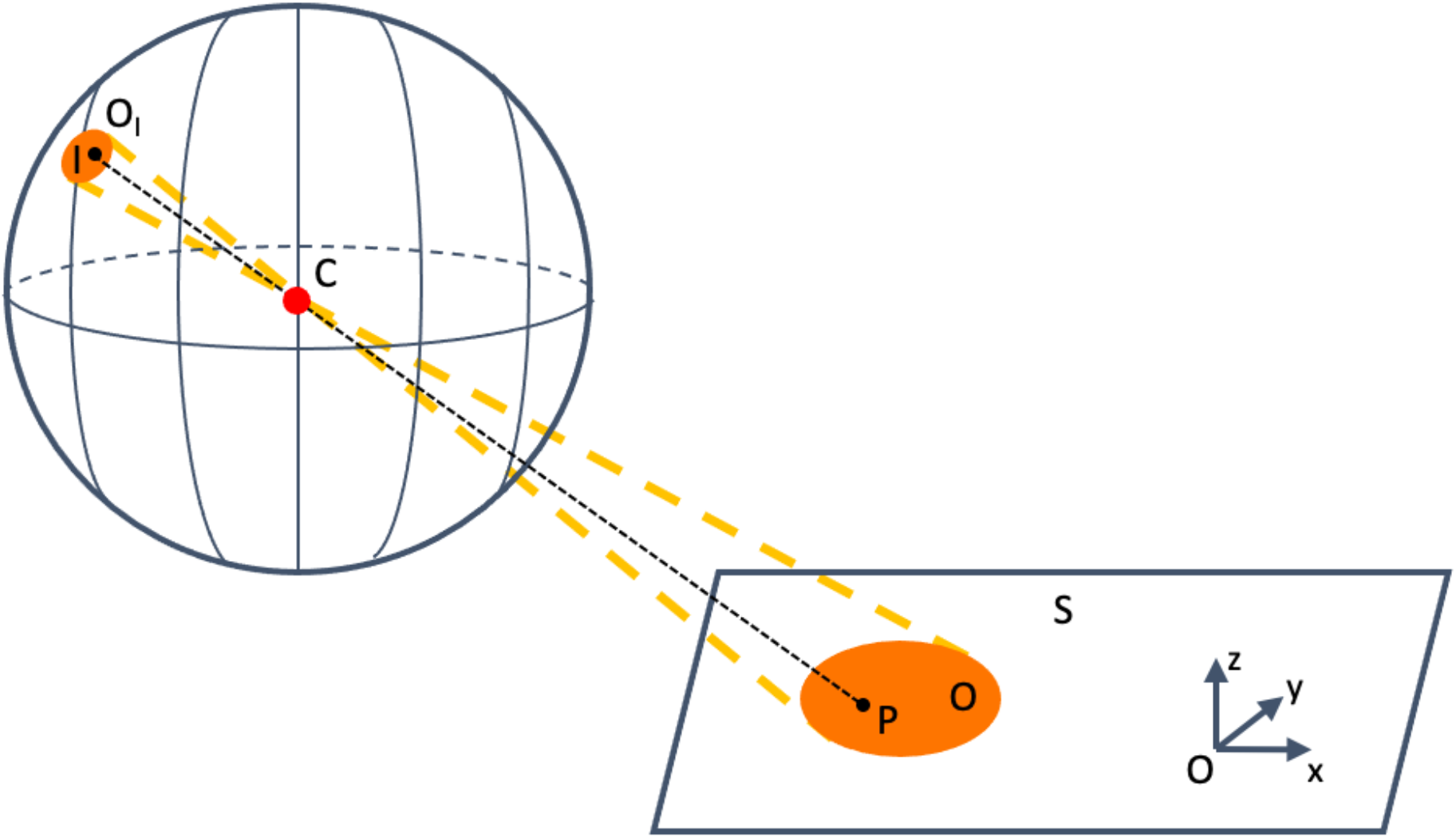

Where S is the plane of the projection screen; O is the projected object onto the screen; P is a point in O; O_I_ is the image of object O on the modeled retina; I is the image of point P on the modeled retina; C is the center of the eye, or the pinhole that allows light into the retina.

The projected 3D position I of point P from the plane S is given by:

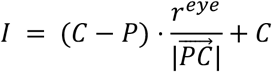

Where *I*,*C*,*P* ∈ *R*^3^, |·| is the euclidean norm and *r^eye^* is the radius of the sphere of the eye. After we trace every point P in O to I, we obtain the resulting image O_I_ on the modeled retina. We then determine the height, width and position of O_I_ on the retina. A toolbox in python implementing this model can be found at: *https://github.com/nguyetming/retina_model*.

Specifically, we used the following parameters when modelling the retina of the larval zebrafish: distance between the eyes: 1.2 mm, eye radius: 0.45 mm, average height of the fish above the projection plane (h): 5 mm, and retina field (see image below): 163°.

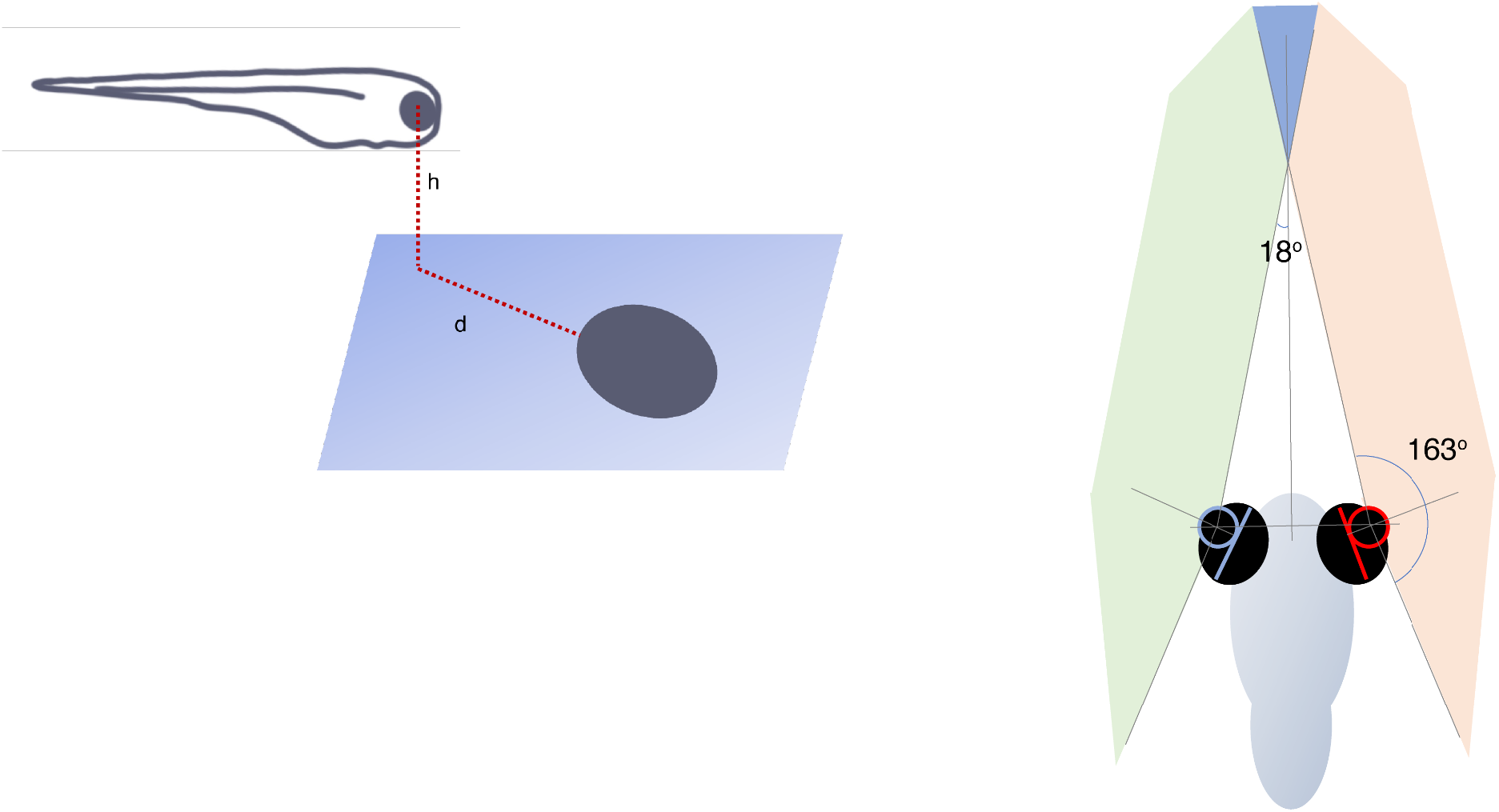

### Modeling groups of free-swimming fish

We simulated groups of N fish ages 7, 14 and 21 dpf, swimming in bounded arenas of sizes 6.5, 9.2 and 12.6 cm interacting according to the algorithms observed in VR experiments (Fig. 4A) or according to the response functions estimated from free swimming experiments (Fig 1F).

a. **Bout size and rate.** In all simulations, each stationary fish, at every time step, probabilistically decides to perform a bout according to the average bout rate observed in group swimming experiments (Fig. S1D). Bout magnitude and bout duration followed that of the average bout calculated from real fish data (Fig. 6SB).
b. **Wall interactions.** When simulated fish were at a distance < 2BL from the walls, they turned away from the wall with probability drawn from the empirical responses of real 7 dpf fish swimming in a group (Fig. S6C). If the executed bout was expected to end outside of the arena, it was truncated to ensure the fish stays inside the simulated arena. When simulated fish were at a distance < 2BL, they did not respond to their neighbors regardless of the model used.
c. **Non-social model.** Simulating N fish that perform wall avoidance at close distances as described above constitutes the non-social model.
d. **Social models based on the clutter integration algorithms extracted from VR.** We used the algorithms we observed in the VR assay, to simulate the interactions between fish in the group according to eq. 1 (see main text) and the clutter response functions extracted from VR experiments (Fig. 4A). In all models, we use the simulated height (H_i_) and distance (d_i_) of each neighbor i to calculate the vertical (V_c_i__) clutter casted on the retina of the focal fish by that neighbor: *V_c_i__* = 2 · *arctan*(*H_i_*/*d_i_*). For simplicity we did not account for occlusions in estimating visual occupancy as initial simulations showed that it did not make a noticeable difference for the group sizes used here. The relative weights *w_i_* assigned to the the responses elicited by visual clutter casted by neighbor i (*p_i_*|*V_i_*)(eq. 1) within each eye followed the weights that best described these responses in the VR experiments (Fig 2E, Fig. 3D): for 7 dpf *w_i_* = *V_i_*/Σ_*i*_ *V_i_* and for 14 and 21 dpf *w_i_* = 1/*N* where N is the number of occupied visual angles (V) in a given eye.
e. **Social models based on algorithms extracted from group swimming experiments.** In these models we used the response functions extracted directly from group swimming experiments to simulate the social interactions of the fish. We calculated the visual angle of each neighbor on the retina of the fish using its width (W_i_), distance (d_i_), and relative orientation (O_i_) to the focal fish. Specifically, we calculated the angle between the vectors pointing from the focal fish to the position of the head and tail of the simulated neighbor. We then summed all visual angles of neighbors within each eye and calculated the difference in occupancy or retinal clutter between the eyes. This value was used to calculate *P*(*turn right*|*Δvisual clutter*) using the inferred response functions from group experiments (Fig. 1F). All other parts of the models are as described in **a-b** above.
f. **Model parameters used in simulations.**

**f. Model parameters used in simulations.**
| Parameter name | Description | Values (7,14,21 dpf) |
| --- | --- | --- |
| Arena diameter | Similar to the arena sizes in group swimming experiments | 6.5, 9.2, 12.6 cm |
| Time interval ($\Delta t$) | Time between simulated steps | 1/50 s for all |
| Simulation time | Total simulated time per group | 600 s for all |
| Number of repetitions | Random repetitions of a given model | 50 for all |
| Fish starting positions | Random positions within 0.9*arena diameter | Same for all |
| Fish length |  | 0.4, 0.5, 0.8 cm |
| Fish height |  | 0.2, 0.25, 0.4 cm |
| Bout Rate | Estimated from group swimming experiments | 1.65, 1.4, 1.4 Hz |
| Bout size | Estimated from group swimming experiments | 0.1 ,0.12 ,0.16 cm |
| Bout duration | Estimated from group swimming experiments | 320 ms |

All modeling codes can be found at:
https://github.com/harpazone/Modeling-larvae-social-behavior

### Sample sizes, trial numbers and power estimation

For all group swimming experiments we used sample sizes that were large to estimate group statistics (e.g. dispersion and alignment) according to previously reported data on collective behavior in zebrafish (20, 29, 53) and to also allow at least 25 degrees of freedom when parametric statistical models were used to compare between experimental conditions. We also chose to test two different group sizes (5 and 10 fish in a group) to show the generality of our findings. In the virtual reality assay, we used 40 trials per stimulus as this number proved sufficient to estimate the response of a single fish to the presented stimuli and 24-32 fish were used per experiment as our preliminary data showed that these are sufficient to estimate the mean responses of fish to the presented stimuli and the differences between these responses for different stimuli.

### Statistical testing

Throughout the paper, we used parametric statistical models to compare experimental conditions, and mean and SEM are reported together with the samples’ data points. Before any model was chosen, we first verified that the underlying assumption of that model, e.g. recommended number of degrees of freedom, homogeneity of variances and the apparent distribution of the data are fulfilled. All reported p-values when comparing two experimental conditions are for the two sided variant of the test.

### Data availability

All data included in this manuscript is available upon request from the corresponding author.

